# Host-derived lipids from tuberculous pleurisy impair macrophage microbicidal-associated metabolic activity

**DOI:** 10.1101/2020.03.23.001818

**Authors:** José Luis Marín Franco, Melanie Genoula, Dan Corral, Gabriel Duette, Malena Ferreyra, Mariano Maio, María Belén Dolotowicz, Omar Emiliano Aparicio-Trejo, Eduardo Patiño-Martínez, Federico Fuentes, Vanessa Soldan, Eduardo José Moraña, Domingo Palmero, Matías Ostrowski, Pablo Schierloh, Carmen Sánchez-Torres, Rogelio Hernández-Pando, José Pedraza-Chaverri, Yoann Rombouts, Emilie Layre, Denis Hudrisier, Christel Vérollet, Olivier Neyrolles, María Del Carmen Sasiain, Geanncarlo Lugo-Villarino, Luciana Balboa

## Abstract

*Mycobacterium tuberculosis* (Mtb) regulates the macrophage metabolic state to thrive in the host. Yet, the responsible mechanisms remain elusive. Macrophage activation towards the microbicidal (M1) program depends on the HIF-1 α-mediated metabolic shift from oxidative phosphorylation towards glycolysis. Here, we asked whether a tuberculosis (TB) microenvironment changes the M1 macrophage metabolic state. We exposed M1 macrophages to the acellular fraction of tuberculous pleural effusions (TB-PE), and found lower glycolytic activity, accompanied by elevated levels of oxidative phosphorylation and bacillary load, compared to controls. The host-derived lipid fraction of TB-PE drove these metabolic alterations. HIF-1α stabilization reverted the effect of TB-PE by restoring M1 metabolism. As a proof-of-concept, Mtb-infected mice with stabilized HIF-1α displayed lower bacillary loads and a pronounced M1-like metabolic profile in alveolar macrophages. Collectively, we demonstrate that host-derived lipids from a TB-associated microenvironment alter the M1 macrophage metabolic reprogramming by hampering HIF-1α functions, thereby impairing control of Mtb infection.

## INTRODUCTION

Tuberculosis (TB) is a highly contagious disease caused by the bacterial pathogen *Mycobacterium tuberculosis* (Mtb). Although treatment of TB is now standardized, it remains one of the top ten causes of death worldwide with 10 million new cases and 1.2 million deaths among human immunodeficiency virus (HIV)-negative people in 2018 (World Health Organization, 2019). Chronic host-pathogen interaction in TB leads to extensive metabolic remodeling in both the host and the pathogen (Russell *et al.*, 2009). Once in the lungs, the alveolar macrophages are the main reservoir of the mycobacteria (Russell *et al.*, 2009). In fact, the success of Mtb as a pathogen largely relies on its ability to adapt to the intracellular milieu of human macrophages. On the one hand, infection leads to activation of host cell-defense programs (*e.g.,* withdrawal of essential nutrients, antimicrobial metabolic pathways) that aim to kill or contain the invading pathogen. On the other hand, the intracellular pathogen can obtain nutrients from the host cell and counteract antimicrobial effector functions (Fuchs *et al.*, 2012; Eisenreich *et al.*, 2013).

Macrophages can modify their metabolic functions from a healing/growth promoting setting (M2 macrophages) towards a killing/inhibitory capacity (M1 macrophages) (Mills *et al.*, 2000; Mills, 2012), representing opposing ends of the full activation spectrum. M1 macrophages, generally induced by interferon (IFN)-γ and/or lipopolysaccharide (LPS) stimulation, are endowed with microbicidal properties; M2 macrophages, generally induced upon interleukin (IL)-4 or IL-13 stimulation, are oriented towards an immunomodulatory and poorly microbicidal phenotype, resulting in impaired anti-mycobacterial properties (Kahnert *et al.*, 2006; Raju *et al.*, 2008; Martinez, Helming and Gordon, 2009). Cellular metabolism and metabolic pathways not only provide energy, but also regulate macrophage phenotype and function (Van den Bossche, O’Neill and Menon, 2017), suggesting that metabolic pathways represent a target for pathogens to evade the host immune response. Particularly, M1 macrophages undergo a metabolic switch towards glycolysis and away from oxidative phosphorylation (OXPHOS) (Kelly and O’Neill, 2015). This glycolytic reprogramming is characterized by an increase in the aerobic glycolysis that results in lactate release, and a decrease in the flux of tricarboxylic acid (TCA) cycle and subsequent OXPHOS (Kelly and O’Neill, 2015; Gleeson *et al.*, 2016). Reduction of the OXPHOS pathway is associated with an increase in the production of mitochondrial reactive oxygen species (mROS) endowed with antimicrobial capacity (West *et al.*, 2011).

A key transcription factor orchestrating the expression of glycolytic enzymes is the hypoxia-inducible factor-1 (HIF-1), a heterodimer comprised of α (HIF-1α) and β (HIF-1β) subunits; HIF-1α being the regulated component of the complex (Wang *et al.*, 1995). The activation of HIF-1α thus results in the production of pro-inflammatory cytokines and other mediators of M1 macrophages, such as glycolytic enzymes and the glucose transporter GLUT1 (Blouin *et al.*, 2004). A different metabolic program is associated with M2 macrophages known to rely on a mixed state driven by enhanced glucose utilization and OXPHOS (Jha *et al.*, 2015; Huang *et al.*, 2016; O’Neill and Pearce, 2016). The M2 program seems to require the induction of fatty acid oxidation (FAO), at least in the murine model (Odegaard and Chawla, 2013; Huang *et al.*, 2014).

Within the framework of macrophage metabolism and its impact on TB, it was recently reported in the mouse model that alveolar macrophages (AMs) and interstitial macrophages (IMs) differ in their ability to control Mtb intracellular growth, which depends on the predominant metabolic status (Huang *et al.*, 2018). A transcriptomic analysis revealed that IMs were glycolytically active, whereas AMs were committed to FAO and exhibited higher susceptibility to bacterial replication than IMs (Huang *et al.*, 2018). Moreover, Mtb infection leads to glycolysis in bone-marrow derived macrophages (Koo, Subbian and Kaplan, 2012; Mehrotra *et al.*, 2014), in lungs of infected mice (Shi *et al.*, 2015), and in lung granulomas from patients with active TB (Subbian *et al.*, 2015). Of note, Mtb infection increases HIF-1 α expression in IFN-γ-activated macrophages, which is essential for the IFN-γ-dependent control of infection (Braverman *et al.*, 2016). Despite these toxic effects enacted by direct infection of cells, Mtb still survives and manages to shift the microenvironment in its favor, which led us to hypothesize that local factors found in the lung milieu may regulate the metabolic network of macrophages and their microbicidal properties. The main goal of our study was to evaluate whether human lung milieu generated during Mtb infection interferes with the metabolic network associated with the M1 macrophage profile and modulates the immune response to favor the mycobacteria.

To address this issue, we used the tuberculous pleural effusion (TB-PE) as a physiologically relevant fluid reflecting the microenvironment found in a human respiratory cavity that is impacted by Mtb infection (Lastrucci *et al.*, 2015; Genoula *et al.*, 2018). The pleural effusion (PE) is an excess of fluid recovered from pleural space characterized by a high protein content and specific leukocytes (Vorster *et al.*, 2015). Although pleural TB is generally categorized as extrapulmonary, there is an intimate anatomic relationship between the pleura and the pulmonary parenchyma (Seibert *et al.*, 1991). The presence of caseating granulomas containing acid-fast bacilli on histological examination of the pleural surface is diagnostic of pleural TB (Gopi *et al.*, 2007), arguing that classic tubercular granulomas can be found in the pleura just like in the lung interstitial tissue. Therefore, TB-PE reflects the microenvironment encountered in the human respiratory cavity during infection, making it a very attractive tool for TB research. In fact, we have employed TB-PE as a physiologically relevant human fluid to study the biology of human monocytes (Balboa *et al.*, 2011; Lastrucci *et al.*, 2015), natural killer cells (Schierloh *et al.*, 2009), B cells (Schierloh *et al.*, 2014; Bénard *et al.*, 2018), gamma delta (γδ) T cells (Yokobori *et al.*, 2009), neutrophils (Alemán *et al.*, 2005) and macrophages (Lugo-Villarino *et al.*, 2018) (including foamy macrophages (Genoula *et al.*, 2018)), and our findings have contributed to the understanding of the impact of the infection on the function of immune components. Moreover, evidence for incomplete control of the infection in pleural TB comes from the fact that patients frequently develop active TB at a later time if untreated (Vorster *et al.*, 2015), suggesting the existence of active mechanisms hampering the protective long-lasting immune response. In this regard, TB-PE shifts *ex vivo* human monocyte differentiation towards an anti-inflammatory M2-like (CD16^+^CD163^+^MerTK^+^pSTAT3^+^) macrophage state, reproducing the phenotype exhibited by macrophages directly isolated from the pleural cavity of TB patients and from lung biopsies of non-human primates with advanced TB (Lastrucci *et al.*, 2015). Likewise, the acellular fraction of TB-PE modifies the lipid metabolism of human macrophages, leading them into foamy macrophages and impacting their effector functions against the bacilli (Genoula *et al.*, 2018).

In this study, we provide strong evidence that TB-PE shifts the glycolysis-based metabolic program of M1 macrophages towards OXPHOS activity by targeting HIF-1α expression. This results in attenuation of the pro-inflammatory and microbicide properties, leading to susceptibility to Mtb intracellular growth in M1 macrophages. Pharmacological stabilization of HIF-1α expression restores not only glycolysis and pro-inflammatory properties in M1 macrophages, but also better control of the Mtb burden in human macrophages and in the lung of Mtb-infected mice. These findings open new avenues for possible host-targeting therapeutic approaches based on HIF-1α modulation, and highlight the need to identify the soluble components in a TB-induced lung milieu responsible for modulation of host cell metabolism.

## RESULTS

### Pleural effusion from active TB patients (TB-PE) inhibits aerobic glycolysis in M1 macrophages

To evaluate whether soluble factors released during Mtb infection interfere with the metabolic network associated with the M1 profile, we generated human macrophages (M0), activated them towards the M1 profile with LPS and IFN-γ in the presence (PE-M1) or absence (M1) of the acellular fraction of tuberculous pleural effusions, and evaluated metabolic parameters as well as M1 macrophage-associated cell-surface markers. TB-PE treatment did not induce cell death in our experimental conditions (Figure S1A-C). The M1-associated cell-surface markers HLA-DR, CD86 and PD-L1 were upregulated to a similar extent in M1 and PE-M1 macrophages in response to IFN-γ and LPS, while CD80 was upregulated to a lesser extent in PE-M1 than M1 macrophages (Figure S1D). Therefore, it appears that the TB-PE treatment does not affect the establishment of the M1 activation program in human macrophages.

Since glucose metabolism in macrophages plays a key role in mounting a robust, pro-inflammatory response to bacterial infection, we evaluated the expression of the glucose transporter GLUT1 (SLC2A1) by flow cytometry using an anti-Glut-1 antibody (Figure 1A) or the Glut-1 ligand H2RBD-EGFP (Kinet *et al.*, 2007) (Figure 1B). We also measured glucose incorporation through the estimation of its consumption from supernatant media (Figure 1C) and the uptake of 2-NBDG (Figure 1D). In line with the literature, M1 macrophages displayed higher levels of both GLUT1 expression and glucose uptake than M0 macrophages, and these levels further increased after TB-PE treatment (Figure 1A-D).

Next, we wondered whether this increased glucose influx was used to fuel the glycolytic pathway leading to the production of lactate. As previously reported (Palsson-Mcdermott *et al.*, 2015), M1 macrophages produced more lactate than M0 cells (Figure 1E). PE-M1 macrophages did not produce significantly more lactate than M0 cells (Figure 1E), whereas those macrophages exposed to PE from other patients *(i.e.,* with cancer, pneumonia or heart failure) did (Figures 1F). We also evaluated the expression of the glycolytic enzymes such as LDHA, which catalyzes the conversion of lactate to pyruvate, and hexokinase 2 (HK2), which produces glucose-6-phosphate and is the predominant isoform in M1 macrophages (Boscá *et al.*, 2015). The upregulation of both enzymes seen in M1 was lower or absent, respectively, in PE-M1 (Figures 1G). Altogether, these results indicate that soluble factors generated by Mtb infection can inhibit the glycolytic metabolic potential of human macrophages during their differentiation towards the microbicidal program.

**figure 1.**
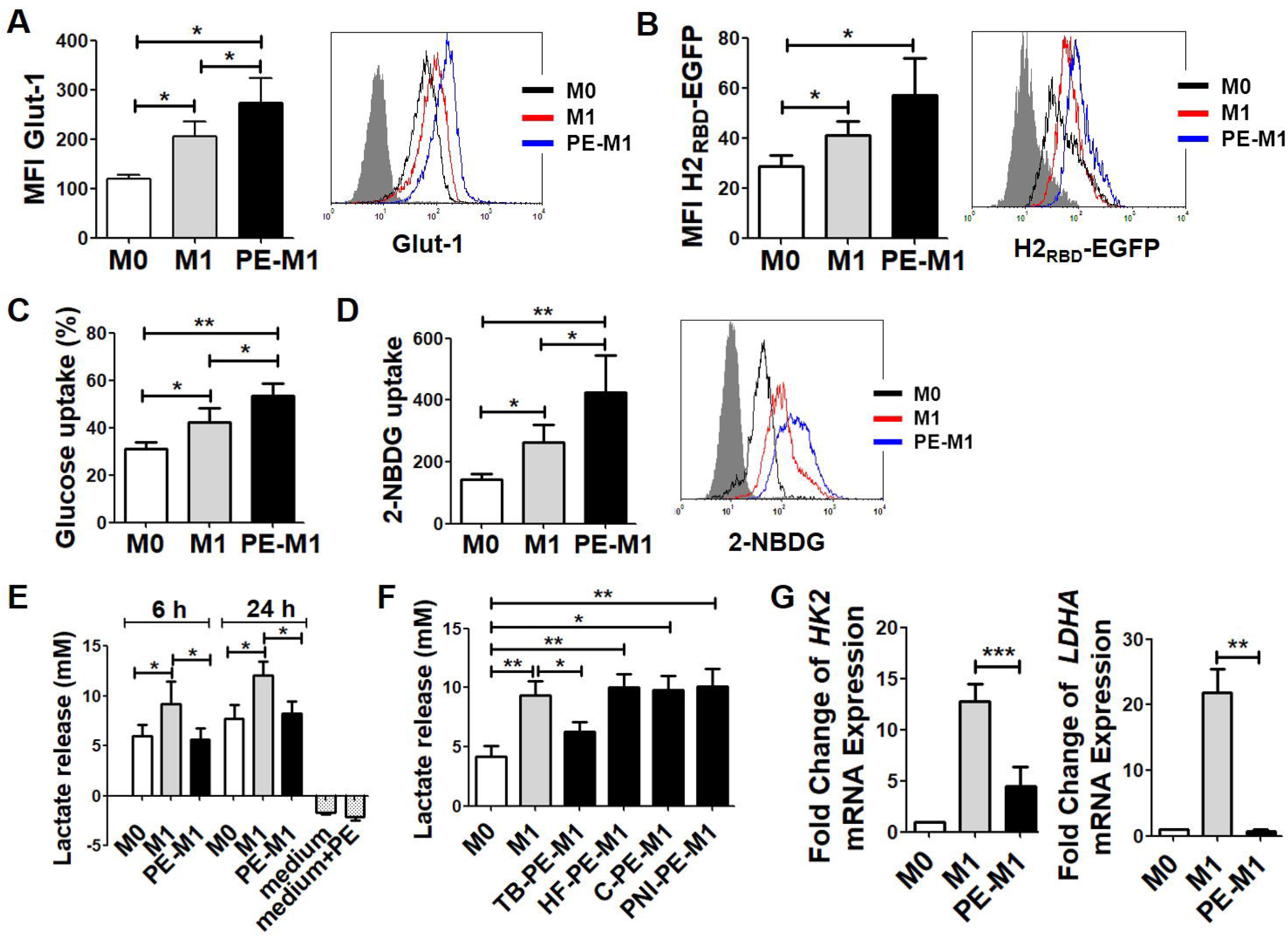
Pleural effusion from active TB patients (TB-PE) inhibits aerobic glycolysis in M1 macrophages. GM-CSF-driven human macrophages (M0) were polarized towards the M1 profile with LPS and IFN-γ in the presence (PE-M1) or absence (M1) of the acellular fraction of tuberculous pleural effusions. (A-B) Mean fluorescence intensity (MFI) of Glut-1 measured though the binding of the cells to the anti-Glut-1 antibody (A, N = 8) and to the Glut-1 ligand H2_RBD_-EGFP (B, N = 4). Representative histograms are shown. (C) Glucose uptake measured in supernatant, N = 6. (D) Glucose consumption measured by 2-NBDG incorporation, N = 5. Representative histogram is shown. (E-F) Lactate release measured in supernatant. HF: heart failure; C: cancer; PNI: parapneumonic PE. N = 8. (G) Expression of lactate dehydrogenase A *(LDHA)* mRNA, and hexokinase 2 *(HK2)* mRNA relative to M0. N = 5. (A-F) Friedman test followed by Dunn’s Multiple Comparison Test: *p < 0.05; **p < 0.01 as depicted by lines. (G) Paired t test. **p < 0.01; ***p < 0.001 for M1 vs PE-M1.

### TB-PE prevents mitochondrial dysfunctions characteristic of M1 macrophages

To further understand the metabolic rewiring induced by TB-PE, we evaluated the mitochondrial function in macrophages by measuring the oxygen consumption rate (OCR) using two different experimental approaches, namely the sensitive high-throughput Seahorse Extracellular Flux Analyzer (Figure 2A-B) and the high-resolution Oxygraph-2k (Figure 2C-D). Both methodologies clearly demonstrated that PE-M1 macrophages have enhanced OXPHOS compared to M1 cells (Figure 2A-D). In fact, PE-M1 macrophages displayed higher basal respiration and ATP-linked OCR (Figure 2A-B), as well as higher OXPHOS-linked OCR and complex IV activity (Figure 2C-D), than M1 macrophages. The decrease in the TCA cycle and electron transport taking place in these cells is known to lead to a reduction in the mitochondrial membrane potential (Palsson-Mcdermott and O’Neill, 2013). Therefore, we assessed the effect of TB-PE on the mitochondrial inner membrane potential. M1 cells displayed a higher percentage of cells with reduced membrane potential than M0 macrophages, while the TB-PE treatment prevented this increase (Figure 2E).

**Figure 2.**
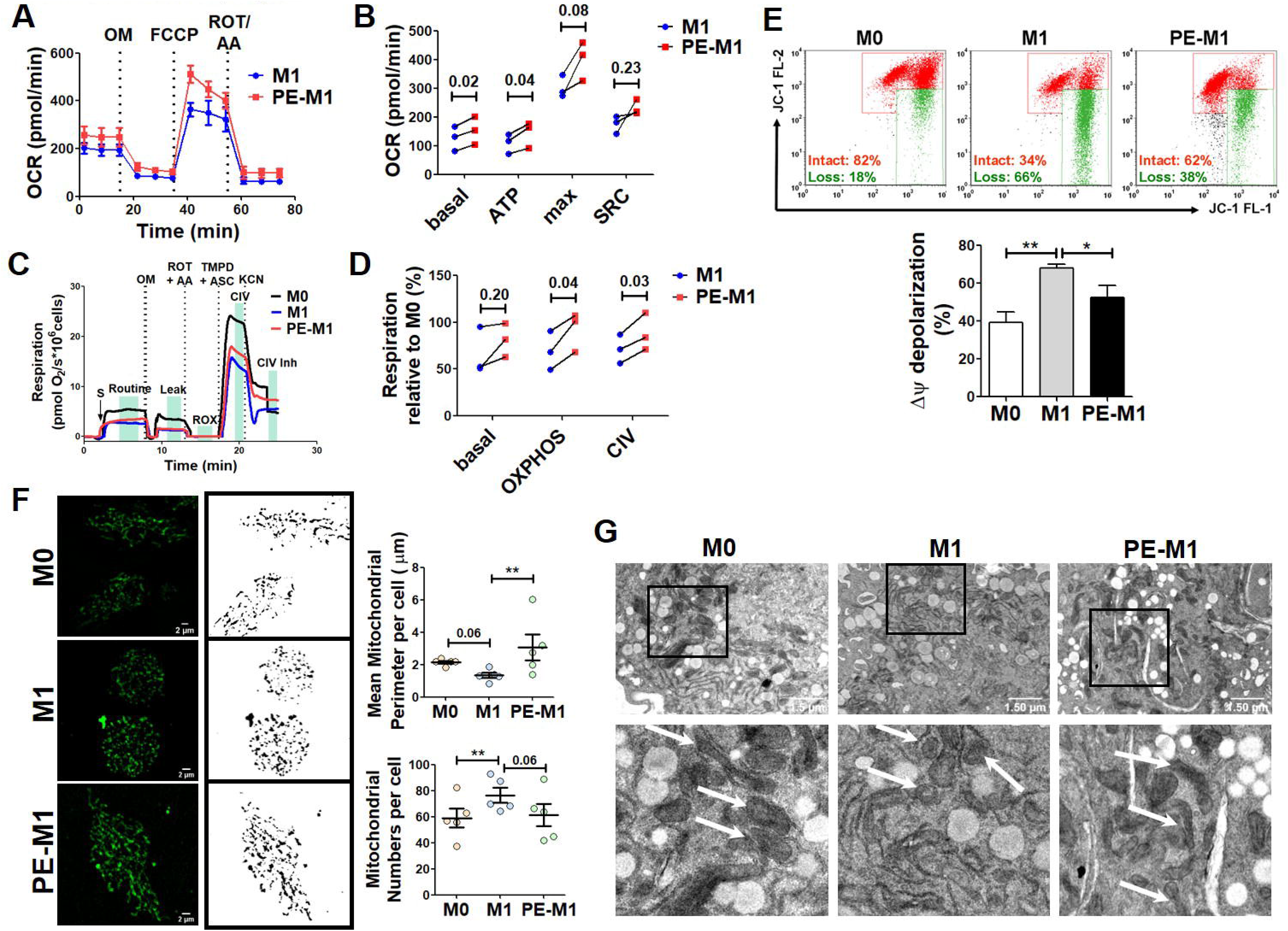
TB-PE prevents mitochondrial dysfunctions characteristic of M1 macrophages. (A-D) OXPHOS parameters assessed by recording the OCR values using a Seahorse Analyzer (A-B) or an Oroboros Oxygraphy-2K (C-D). (A and C) One representative experiment (triplicates). (B and D) Respiratory parameters measured in 3 independent experiments. Student’s T-test. (E) Mitochondrial membrane potential measured by JC-1 probe staining, N=6. (F) Analyses of mitochondrial morphology by confocal microscopy in macrophages. Representative Z-stacks of images of Green Mitospy-stained macrophages and their corresponding grayscale mask image. Right panels show the mean mitochondrial perimeter and numbers of mitochondria per cell from 5 donors. (G) Representative electron microscopy micrographs of macrophages showing mitochondrial ultrastructure. White arrows denote mitochondria. (B and D) Paired t test. (E-F) Friedman test followed by Dunn’s Multiple Comparison Test: *p < 0.05; **p < 0.01 as depicted by lines.

Mitochondrial shape and bioenergetics are intimately linked(Rambold and Pearce, 2018), and mitochondrial morphology is altered in macrophages stimulated with LPS leading to small, diverse mitochondria dispersed throughout the cytoplasm (Gao *et al.*, 2017). Confocal microscopy of macrophages labeled with Mitospy, which stains mitochondria regardless of their membrane potential, revealed punctate and more numerous mitochondria in M1 macrophages, while in PE-M1 cells the organelles were more tubular-shaped and less numerous, similar to M0 cells (Figure 2F). Transmission electron microscopy confirmed that, compared to M0 cells, mitochondria in M1 macrophages were smaller and displayed a disrupted cristae shape, while these defects were not seen in PE-M1 cells (Figure 2G). These findings support the hypothesis that soluble factors generated during Mtb infection block the shift towards the aerobic glycolysis that is usually accompanied by mitochondrial dysfunction during the activation of microbicidal macrophages.

### TB-PE inhibits HIF-1α expression and pro-inflammatory functions in M1 macrophages

To address potential mechanisms underlying the inhibitory effects of TB-PE, we evaluated the expression of HIF-1α, since this transcription factor governs the metabolic reprogramming by increasing the expression of glycolytic enzymes and pro-inflammatory cytokines *(e.g.,* IL-1β), while inhibiting OXPHOS activity (Palsson-Mcdermott *et al.*, 2015; Gleeson and Sheedy, 2016). Indeed, HIF-1α was induced in M1 cells, but not in PE-M1 cells, at the protein and RNA level (Figure 3A-B). An important enhancer of HIF-1α activity is the mammalian target of rapamycin (mTOR) complex 1 (mTORC1). HIF-1α can be activated by mTORC1 upon cytokine signaling, by elevated levels of succinate, or by reactive oxygen species (ROS) (Finlay *et al.*, 2012; Tannahill *et al.*, 2013; Movafagh, Crook and Vo, 2015). Upon activation, mTOR is phosphorylated on several residues including S2448; while we found that this phosphorylated form was upregulated in M1, we failed to detect it in PE-M1 cells (Figure 3C). Bacterial infection, or LPS stimulation, results in increased NF-κB activity in phagocytes, which in turn induces *HIF-1α* mRNA transcription (Rius *et al.*, 2008). Pro-inflammatory features such as the expression of NF-κB, the release of IL-1β and the production of mROS were less prominent in PE-M1 than M1 cells (Figure 3D-F). Hence, the soluble factors generated by Mtb infection exert their inhibitory effect on pro-inflammatory functions as well as on HIF-1α activity.

**Figure 3.**
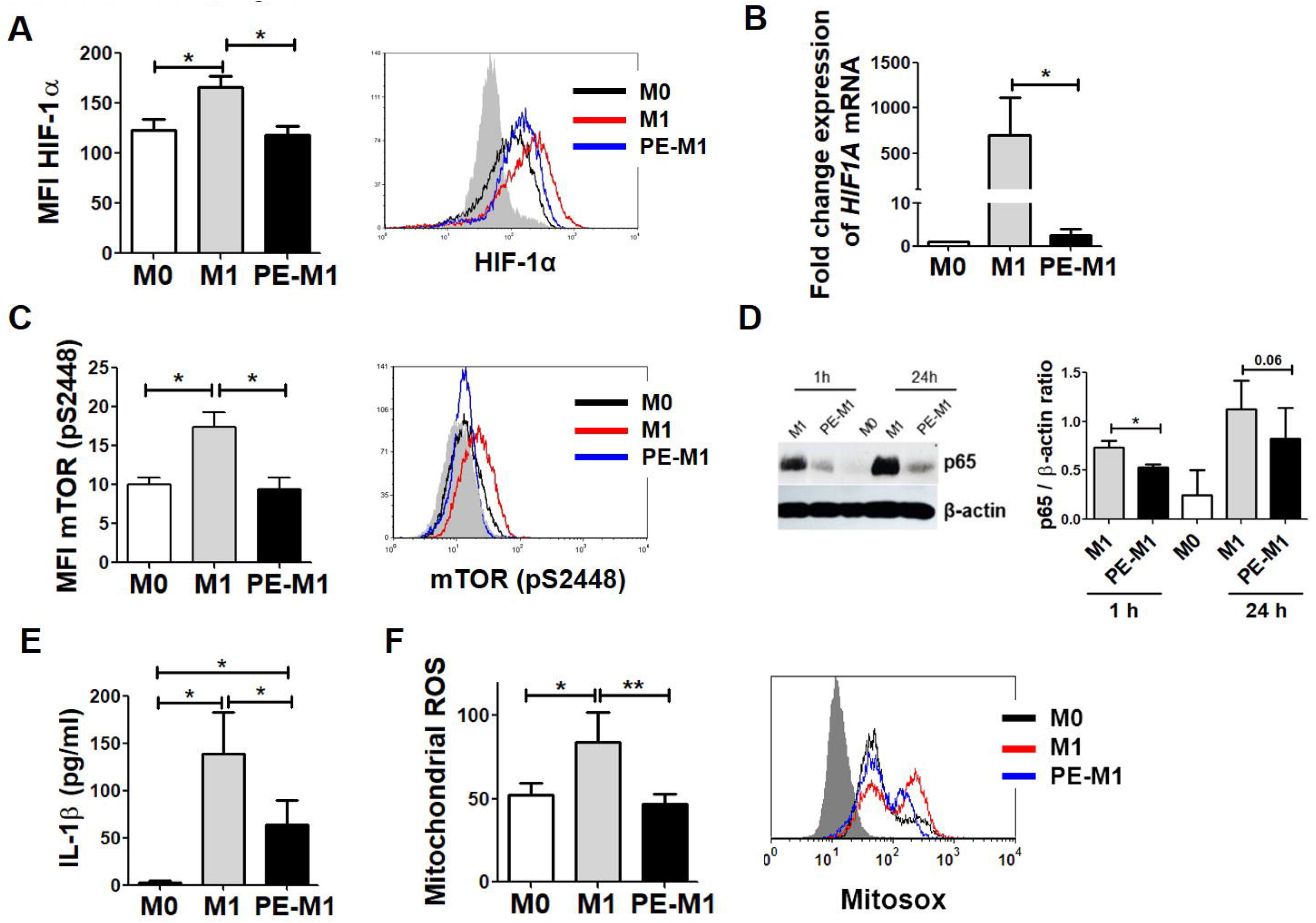
TB-PE inhibits HIF-1α expression and pro-inflammatory functions in M1 macrophages. (A) Left: mean fluorescence intensity (MFI) of HIF-1α measured by FACS, N = 7. Right: representative histogram. (B) Expression of *HIF-1α* mRNA relative to M0, N = 6. (C) Left: percentages of mTOR (pS2448) positive cells measured by FACS, N = 5. Right: representative histogram. (D) Analysis of p65 and β-actin protein expression level at 1 and 24 h post-stimulation by western Blot (left panel) and quantification (right panel; N = 4) macrophages. (E) IL-1β production by ELISA, N = 4. (F) Left: MFI quantification of MitoSOX for human macrophages, N=12. Right: representative histogram. Friedman test followed by Dunn’s Multiple Comparison Test: *p < 0.05; **p < 0.01 as depicted by lines.

### TB-PE inhibits pro-inflammatory functions in human macrophages in response to Mtb

Mtb infection leads to glycolysis in bone marrow-derived macrophages (Koo, Subbian and Kaplan, 2012; Mehrotra *et al.*, 2014) and to increased HIF-1α expression in IFN-γ-activated macrophages (Braverman *et al.*, 2016). We confirmed this is also the case in IFN-γ-polarized macrophages stimulated with irradiated Mtb or infected with live Mtb, since both lactate release and HIF-1α expression increased (Figure 4A-D). Different strains of Mtb displayed similar capacity to induce the metabolic reprogramming in IFN-γ-polarized macrophages, as measured by lactate release and glucose consumption (Figure S2A-B). TB-PE treatment reduced the release of lactate and the expression of HIF-1α driven by irradiated or live Mtb (Figure 4A-D). Furthermore, HIF-1α-associated production of mROS and IL-1β in response to irradiated Mtb was reduced in PE-treated macrophages (Figure 4E-F), and these cells showed an increase in live Mtb load reflecting their lower microbicidal activity (Figure 4G). Concerning the surface marker expression, macrophages stimulated with IFN-γ and irradiated Mtb up-regulated the levels of HLA-DR, CD86, PD-L1, and CD80 regardless of TB-PE treatment (Figure S2C-D). These observations establish that, beyond the classical activation of macrophages with IFN-γ/LPS, soluble factors generated by Mtb infection can also inhibit the activation of pro-inflammatory and metabolic properties in these cells interacting directly with the pathogen.

**Figure 4.**
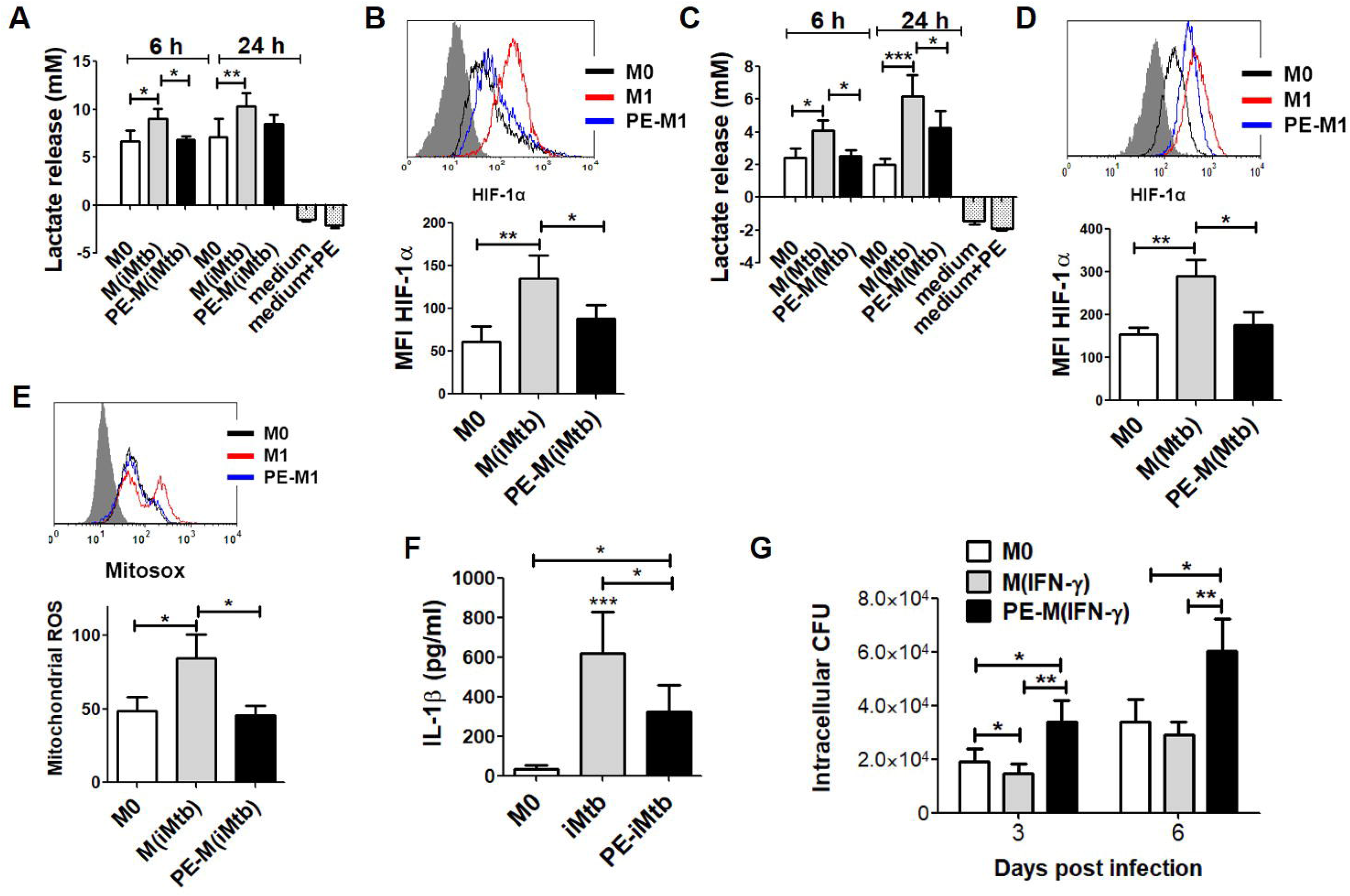
TB-PE inhibits pro-inflammatory functions in human macrophages in response to Mtb. GM-CSF-driven human macrophages (M0) were stimulated with IFN-γ and irradiated Mtb or live Mtb in the presence (PE-M(iMtb) and PE-M(Mtb)) or absence (M(iMtb) and M(Mtb)) of the acellular fraction of tuberculous pleural effusions. (A and C) Lactate release by macrophages in responses to stimulation or not with irradiated (A) or live (C) Mtb in the presence or absence of PE at 6 and 24 h, N=8 (A) and N=9 (C). (B and D) Mean fluorescence intensity (MFI) of HIF-1α induced by irradiated (B) or live (D) Mtb at 24 h, N = 7 (B) and N = 8 (D). (E) MFI quantification of MitoSOX for human macrophages in responses to stimulation or not with irradiated Mtb H37Rv in the presence or absence of PE at 24 h, N = 5. (F) IL-1β cytokine levels produced by macrophages in responses to stimulation or not with irradiated Mtb H37Rv in the presence or absence of PE at 24 h, N = 5. (G) Intracellular colony forming units (CFU) were determined at different time points in macrophages (M0) stimulated with IFN-γ in the presence (PE-M(IFN-γ)) or absence (M(IFN-γ)) of PE for 24 h, washed and infected with Mtb, N = 7. Friedman test followed by Dunn’s Multiple Comparison Test: *p < 0.05; **p < 0.01; ***p < 0.001 as depicted by lines.

### HIF-1α stabilization reverts the inhibitory effects of TB-PE on the pro-inflammatory macrophage program

To establish the role of HIF-1α as the target of the TB-PE treatment, we stabilized its activity by blocking the HIF prolyl hydroxylases (Jaakkola *et al.*, 2001). To this end, we used inhibitors of prolyl hydroxylases, such as dimethyloxalylglycine (DMOG) and Cobalt (II) chloride (CoCl_2_). HIF-1α stabilization in PE-M1 cells treated with DMOG or CoCl_2_ resulted in higher expression of HIF-1α, HK2 and LDHA, as well as an increase in lactate production and in mitochondrial fragmentation and cristae defects (Figure 5A-E). Of note, *LDHA* and *HK2* are target genes under the transcriptional control of HIF-1α (Boscá *et al.*, 2015; Semba *et al.*, 2016). These phenotypic changes induced by HIF-1α stabilization were associated with the recovery of pro-inflammatory functions, such as IL-1β and mROS production (Figure 5F-G). Along with these functional changes, TB-PE also promoted a poor capacity to control the bacillary load in IFN-γ-activated macrophages, a feature that was reverted by DMOG treatment (Figure 5H-I). These data establish a central role for the inhibition of HIF-1α activity in the impaired microbicidal activity of macrophages triggered by soluble factors generated during Mtb infection.

**Figure 5.**
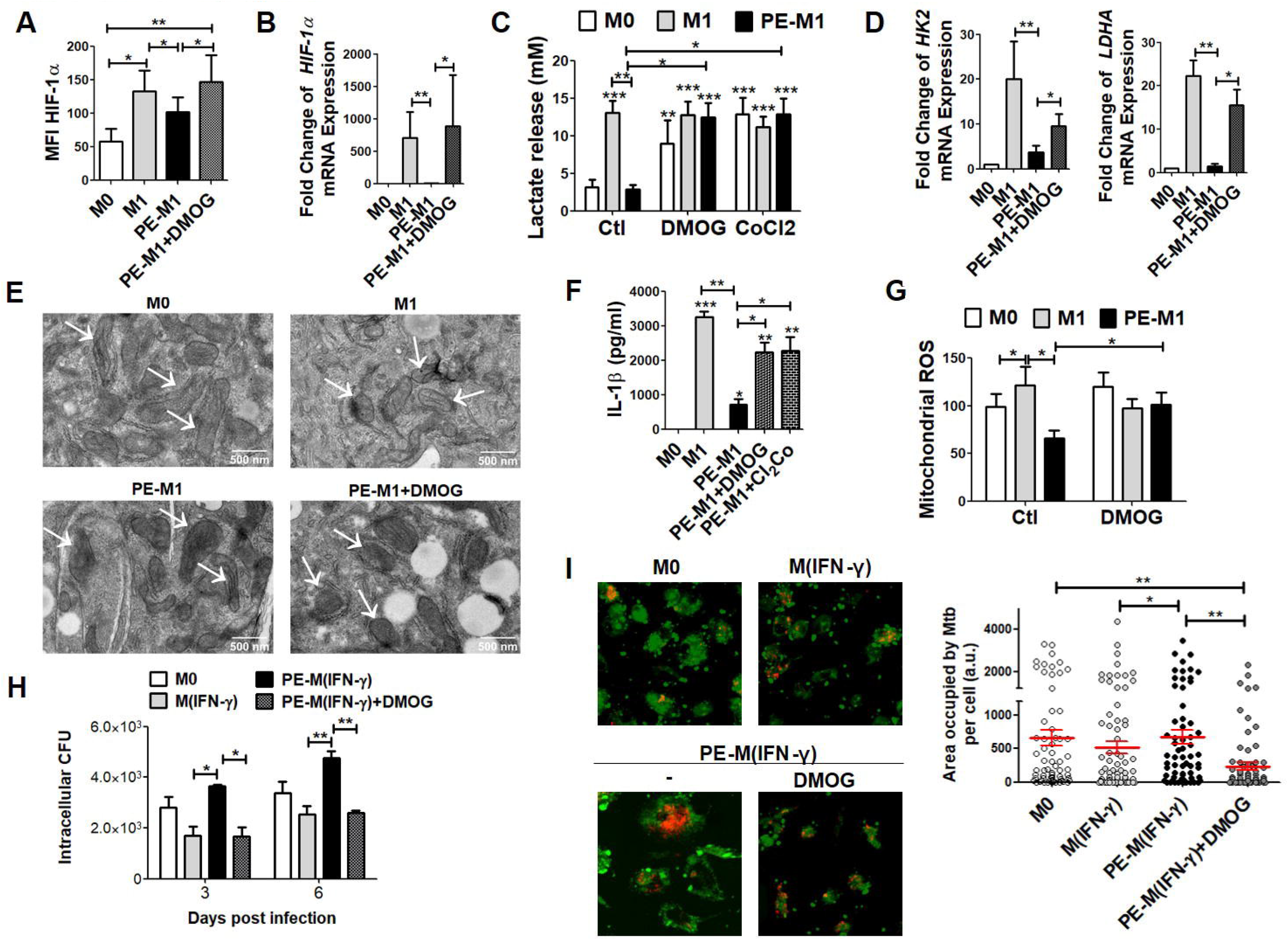
HIF-1α stabilization reverts the inhibitory effects of TB-PE on the pro-inflammatory macrophage program. GM-CSF-driven human macrophages (M0) were polarized towards the M1 profile with LPS and IFN-γ in the presence (PE-M1) or absence (M1) of the acellular fraction of tuberculous pleural effusions with or without DMOG or CoCl_2_. (A) HIF-1α expression in macrophages, N = 7. (B) Expression of *HIF-1α* mRNA relative to M0, N = 6. (C) Lactate release by macrophages, N = 6. (D) Expression of lactate dehydrogenase A *(LDHA)* mRNA and hexokinase 2 *(HK2)* mRNA relative to M0, N = 7. (E) Representative electron microscopy micrographs of macrophages showing mitochondrial ultrastructure. White arrows denote mitochondria. (F) IL-1β production by ELISA, N = 7. (G) MFI quantification of MitoSOX for human macrophages, N = 7. (H) M0 and IFN-γ-activated macrophages were infected with Mtb and exposed or not to TB-PE and DMOG. Intracellular colony forming units were determined at different time points, N = 4. Friedman test followed by Dunn’s Multiple Comparison Test: *p < 0.05; **p < 0.01; as depicted by lines. (I) M0 and IFN-γ-activated macrophages were infected with RFP-expressing Mtb and exposed or not to TB-PE and DMOG. The area occupied by Mtb per cell was assessed in z-stacks from confocal laser scanning microscopy images at 48 post-infection. One representative experiment showing 80-100 cells per condition is shown. ONE-way ANOVA followed by Bonferroni’s Multiple Comparison Test: *p < 0.05; **p < 0.01; ***p < 0.001 as depicted by lines.

### HIF-1α stabilization improves mycobacterial control in mice and is associated with mitochondrial stress in alveolar macrophages

Next, we evaluated the effect of DMOG on the bacillary loads and on the mitochondrial functions of pulmonary macrophages from Mtb-infected mice treated (or not) with DMOG at early stages of Mtb infection. DMOG-treated mice had lower pulmonary bacillary loads at 14 days post-infection than control mice (Figure 6A). We then followed a flow cytometric gating protocol based on previous analysis to distinguish different types of mononuclear phagocytes present in lungs of Mtb-infected mice (Huang *et al.*, 2018) (Figure S3 A). DMOG-treated mice had a higher percentage (but not a higher absolute number) of AMs than control mice (Figure 6B-C). No significant differences were found in the percentage or absolute number of IMs (Figure 6D-E), or lung polymorphonuclear leukocytes (Figure S3B-C). In addition, both AMs and IMs from DMOG-treated animals had lower mitochondrial inner membrane potential (Figure 6F-H and Figure S3D-F) and higher mROS production than control AMs (Figure 6I-K and Figure S3G-I). These results indicate that DMOG treatment can protect against Mtb infection, and this protection is associated with an increase in AM mitochondrial stress, suggesting a metabolic switch.

**Figure 6.**
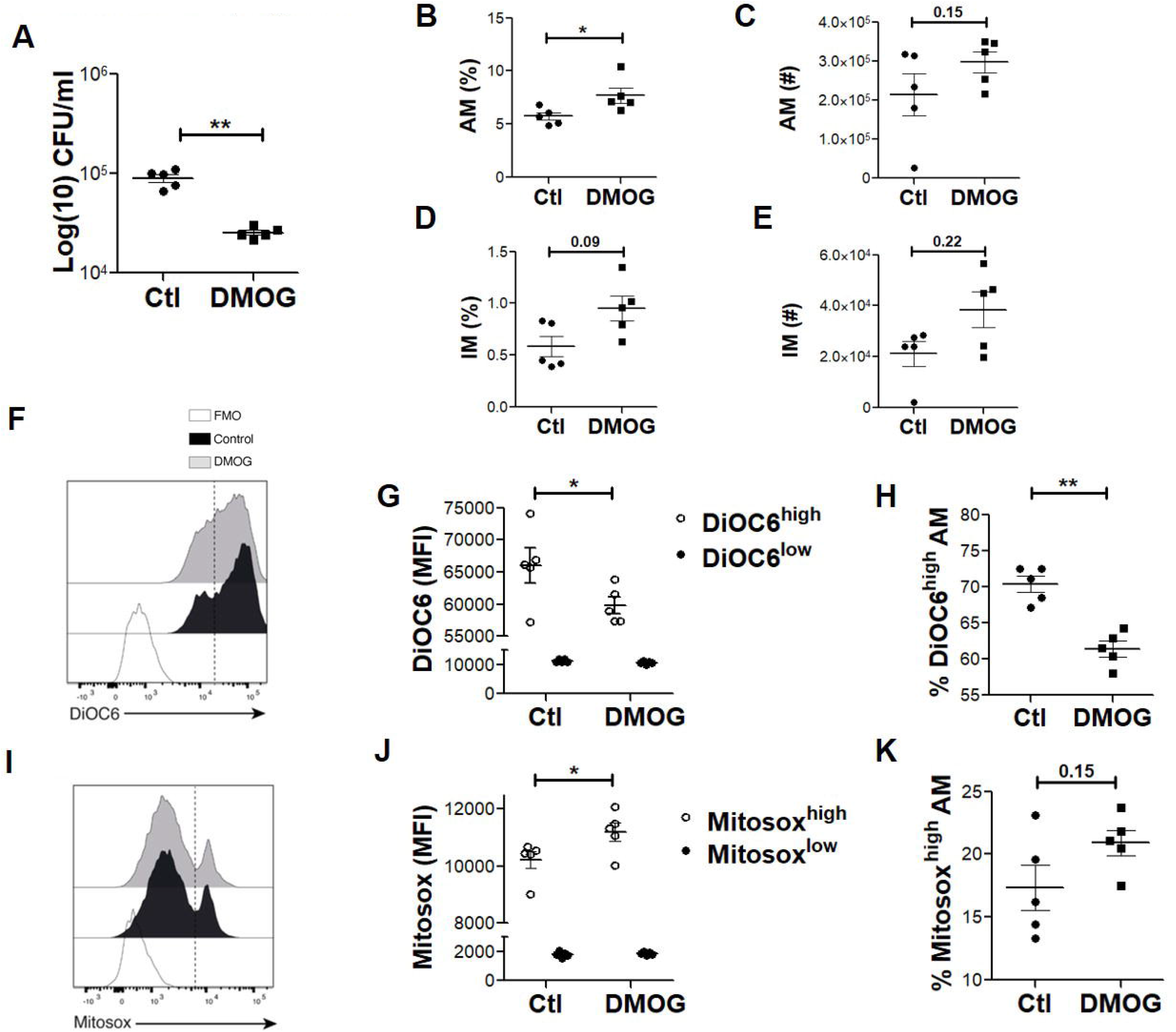
HIF-1α stabilization improves mycobacterial control in mice and is associated with mitochondrial stress in alveolar macrophages. Mice were infected intranasally with Mtb (1000 UFC per mouse), and they were treated or not with DMOG for 14 days. (A) Colony forming units (CFU) recovered from lungs of DMOG-treated mice or control mice (Ctl). A representative experiment of three independent experiments is shown. (B-C) Percentages (%) of alveolar macrophages (AM) among CD45.2^+^ Live cells (B) and absolute number (#) of AM (C). A representative of two independent experiments is shown. (D-E) Percentages of interstitial macrophages (IM) among CD45.2^+^ Live cells (D) and absolute number of AM (E). A representative experiment of two independent experiments is shown. (F) Representative histogram of DiOC6 labeling for AM from DMOG-treated mice, control mice and fluorescent minus one (FMO). Vertical dashed line separates high vs low events. (G) Mean fluorescent intensity (MFI) quantification of DiOC6 for AM (H) Percentages of DiOC6^high^ AM. A representative of two independent experiments is shown. (I) Representative histogram of MitoSOX labeling for AM from DMOG-treated mice, control mice and FMO. (J) MFI quantification of MitoSOX for AM. Vertical dashed line separates high vs low events. (K) Percentages of MitoSOX^hlgh^ AM. A representative of two independent experiments is shown. Mann-Whitney test: *p < 0,05; **p< 0,01; Two-way ANOVA: *p < 0,05.

### Host-derived lipids from TB-PE are responsible for the metabolic remodeling activity in M1 macrophages

To investigate the nature of the TB-PE components responsible for the metabolic alterations observed in human macrophages, we tested the metabolic activity of the TB-PE fraction obtained after treatment with proteinase K, which was embedded onto polyacrylamide hydrogel (Pk-PE). Protein digestion was controlled using SDS-PAGE, followed by Coomassie blue staining (Figure S4A). TB-PE digested of proteins was still able to diminish the release of lactate driven by IFN-γ/LPS without inducing cell death in M1 macrophages (Figure S4B-C). Thereafter, we separated two fractions of TB-PE as follows: one enriched in polar metabolites (PMPE) and the other comprising lipids (LPE), and used them to treat M1 macrophages at the same concentration as in the untouched TB-PE (1x) or at a double dose (2x). LPE but not PMPE was able to reduce the release of lactate at comparable doses found in intact TB-PE (Figure 7A). Cell viability was not modified either by PMPE or LPE treatment at 1x or 2x doses (Figure 7B and S4D). In line with the reduction of lactate, LPE also inhibited HIF-1α expression (Figure 7C), partially decreased mROS production by M1 macrophages (Figure 7D), and rendered the IFN-γ-activated macrophages more susceptible to Mtb intracellular growth (Figure 7E).

**Figure 7.**
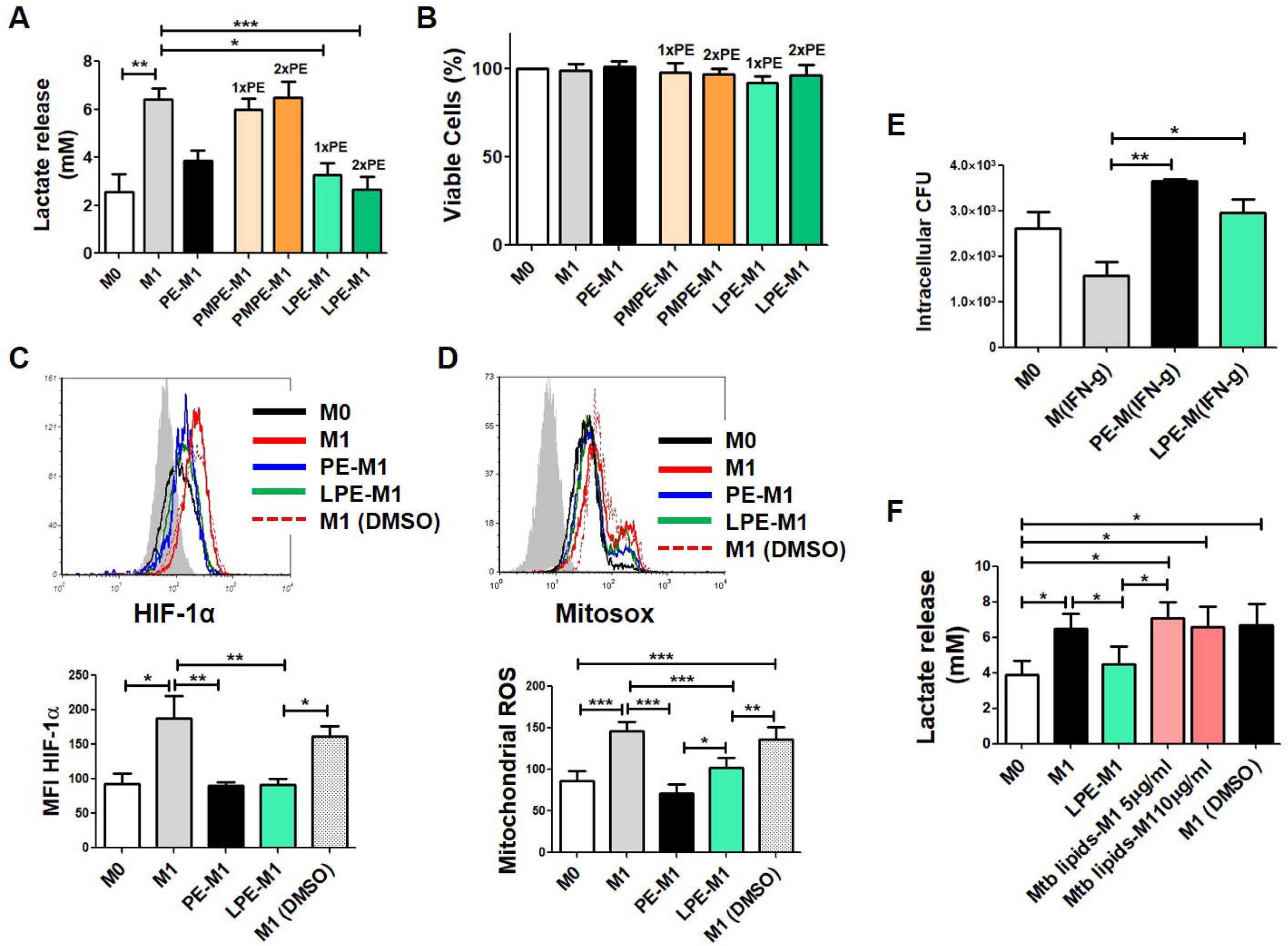
Host-derived lipids from TB-PE are responsible for the metabolic remodeling activity in M1 macrophages. (A-B) Lactate release (A) and cell viability (B) of M0, M1, PE-M1, PMPE-M1 (Polar metabolites PE-M1) or LPE-M1 (Lipids PE-M1) cells, N = 8. PMPE and LPE fractions were used at the same concentration as in the intact TB-PE (1x) or at a double dose (2x). (C) HIF-1α expression in macrophages, N = 6. (D) MFI quantification of MitoSOX for macrophages, N = 8. (E) M0 and IFN-γ-activated macrophages were infected with Mtb and exposed or not to TB-PE or LPE. Intracellular colony forming units determined at day 3 post-infection, N = 4. (F) Lactate release by macrophages exposed or not to total lipids fraction obtained from Mtb, N = 6. Friedman test followed by Dunn’s Multiple Comparison Test: *p < 0.05; **p < 0.01; ***p < 0.001, as depicted by lines.

To determine whether metabolic-active lipids found in TB-PE were bacterial or host-derived, we performed a lipidomic analysis of the LPE. Mass spectrometry signals corresponding to the main mycobacterial lipid families were not detected in the TB-PE lipid extract dataset, according to MycoMass and My coMap databases (Layre *et al.*, 2011), supporting the idea that the activity of TB-PE lipid extract is likely to be driven by host lipids. This notion was further supported by the fact that the total lipid fraction obtained from Mtb was not able to impair the release of lactate by M1 macrophages in our experimental settings (Figure 7F and S4E). These results demonstrated that host-derived lipid components elicited by Mtb infection are responsible for rewiring the metabolism of M1 macrophages impacting the control of the bacillary intracellular load.

## DISCUSSION

TB, as a chronic condition, entails the establishment of extensive metabolic remodeling in both host and pathogen (Russell *et al.*, 2010). Our ability to control the burden of TB globally is limited by the absence of an effective vaccine, an aspect which is worsened by the lack of any reliable immune correlates of local tissue protective immunity (Bhatt *et al.*, 2015; Goletti *et al.*, 2016; Huang and Russell, 2017). Progress in these areas requires a more profound understanding of the biology of Mtb and its interaction with the human host and the implementation of appropriate experimental models to investigate TB (Karp, Wilson and Stuart, 2015). Therefore, more basic research focused on human physiologically relevant samples derived from active TB patients is required to perform functional and mechanistic studies on TB under physiological conditions, especially those samples that reflect the complex environment of human lung tissue. In this regard, numerous studies have focused on the investigation of the broncho-alveolar lavage and, relatively less, lung tissue explants obtained from patients undergoing lung surgery for clinical reasons unrelated to pulmonary infection, which were infected *in vitro* with Mtb (Ganbat *et al.*, 2016; Maertzdorf *et al.*, 2018). In the former case, to obtain the broncho-alveolar lavage, a saline solution must be instilled in the lung causing the unavoidable dilution of the lung milieu. In addition, broncho-alveolar lavages usually result in an incomplete sample and poor representative of the lower lung airways; it is also an invasive and risky procedure, making subjects less reluctant to participate. In the latter case, there is no leukocyte recruitment to the site of infection, making these lung tissue explants less representative of the lung microenvironment generated in the present of leukocytes. In comparison, as a therapeutic procedure to drain the effusion for clinical diagnosis that also improves patient clinical conditions, we argue that PE is an unperturbed microenvironment generated by Mtb infection and contains most of the leukocyte populations that are involved in TB immunity, including monocytes and macrophages. Although we are aware that *ex vivo/in vitro* studies do not fully reflect the complexity of the lung architecture and its impact on host–pathogen interactions, the use of the acellular fraction of the infection site provides a unique tool to study how a *bona fide* TB-associated milieu influences the host cells. In this regard, we have previously used the acellular fraction of TB-PE and demonstrated its pertinence as a model human lung TB microenvironment (Lastrucci *et al.*, 2015; Genoula *et al.*, 2018; Souriant *et al.*, 2018).

Here, we show that host-derived lipids present within the acellular fraction of TB-PE alter the metabolic reprogramming of M1 macrophages by targeting HIF-1α expression. In our experimental model, we exposed GM-CSF-driven human macrophages to M1-activating stimuli in the presence (or not) of the TB-PE to reproduce the complex milieu in which the activation program may take place physiologically. Even though GM-CSF itself skews macrophages towards glycolysis (Na *et al.*, 2017), we considered these as “M0” cells to emphasize their lower activation state in comparison to those stimulated with LPS or Mtb. Yet, these cells are very different from the classical “M0” generated under M-CSF treatment (Jaguin *et al.*, 2013). In terms of metabolic parameters, we demonstrated that even when PE-M1 macrophages showed a higher consumption of glucose, this additional glucose is not converted into lactate as in M1 cells, but rather used to feed the OXPHOS pathway, adopting an M2-like metabolic profile. In this regard, although it was widely accepted that a higher consumption of glucose associates with M1 macrophages leading to a higher aerobic glycolytic activity, it is now known that M2 macrophages *(e.g.,* driven by IL-4) also exhibit enhanced glucose utilization during their activation program; higher glucose uptake is used to feed the mitochondrial respiration (Huang *et al.*, 2016). In line with this, TB-PE-M1 macrophages, shown to be more prone to OXPHOS than M1 cells, also displayed an enhanced consumption of glucose. The less glycolytic profile established in PE-M1 macrophages is further supported by the reduced levels of lactate and the decreased expression of LDHA and HK2, both HIF-1α target genes (Boscá *et al.*, 2015; Semba *et al.*, 2016). Additionally, we demonstrated that TB-PE impairs M1-associated metabolism, but it also leads to a mixed repertoire of cell-surface markers in macrophages, reflecting the complex nature of TB-PE as a physiologically relevant fluid.

Another line of supporting evidence for the capacity of TB-PE to influence the metabolic state of microbicidal macrophages is the change in mitochondrial morphology that is compatible with a more active oxidative metabolism, such as: 1) tubular shaped mitochondria and 2) functional cristae with folding architecture of the mitochondrial inner membrane. The fusion process between mitochondria is important for OXPHOS activity, while the fission process leads to a higher number of smaller and punctuated organelles, which have an impaired mitochondrial respiration activity (Rambold and Pearce, 2018). In this regard, LPS-stimulated macrophages have small, diverse mitochondria dispersed throughout the cytoplasm (Gao *et al.*, 2017), which we confirmed in our IFN-γ/LPS-stimulated macrophages. This mitochondrial fragmentation observed in M1 macrophages is not achieved when cells are exposed to TB-PE. Also, changes in the ultrastructure of mitochondria are associated with the regulation of mitochondrial functions; that is, mitochondrial cristae are the main site of OXPHOS and their folded structure is critical for respiratory chain supercomplex assembly (Cogliati, Enriquez and Scorrano, 2016). In this context, we showed a normalization of mitochondrial cristae morphology in PE-M1 cells compared with M1 macrophages, which is consistent with the elevated OXPHOS activity found in PE-M1 cells. Particularly, the loss of mitochondrial membrane potential is associated with accumulation of mROS (Cai *et al.*, 2015; Weinberg, Sena and Chandel, 2015; Ip *et al.*, 2017). Also, mitochondrial depolarization is central to the increased microbicidal ability of macrophages under hypoxia or upon activation in response to Mtb infection (Matta and Kumar, 2016). In agreement with these findings, PE-M1 macrophages exhibited neither loss of mitochondrial membrane potential nor an appreciable induction of mROS upon LPS stimulation, displaying impaired ability to control Mtb intracellular load. Metabolic repurposing of mitochondria away from OXPHOS is key to the regulation of pro-inflammatory responses, including the expression of pro-IL-1β and the generation of mROS via reverse electron transport (Van den Bossche, O’Neill and Menon, 2017). Indeed, the pro-inflammatory cytokine IL-1β is a crucial mediator of inflammation and plays a vital role in the host resistance to Mtb infections (Juffermans *et al.*, 2000). In the case of mROS, it is thought that professional phagocytes generate ROS primarily via the phagosomal nicotinamide adenine dinucleotide phosphate (NADPH) oxidase machinery, contributing to macrophage bactericidal activity (West *et al.*, 2011). mROS can be delivered to *Staphylococcus aureus*-containing phagosomes via mitochondria-derived vesicles (Abuaita, Schultz and O’Riordan, 2018), providing a putative explanation for how mROS can reach bacteria within the phagosomal lumen. Regardless of the direct microbicidal effect of mROS, these species can also contribute to the pro-inflammatory profile by promoting the expression of HIF-1α (Wang, Malo and Hekimi, 2010). Incidentally, it was recently reported that the glycolysis mediated by HIF-1α is a basic defense mechanism of macrophages against Mtb infection (Osada-Oka *et al.*, 2019). The proposed antimycobacterial mechanism is based on the notion that pyruvate may be more efficiently used than glucose as the carbon source for intracellular growth of Mtb within the phagosome. HIF-1α activation has a protective mechanism by hijacking pyruvate levels as a consequence of fostering its conversion into lactate through LDHA up-regulation (Osada-Oka *et al.*, 2019).

HIF-1α and mTOR are now recognized as the key regulators of this shift of metabolism toward glycolysis (Srivastava and Mannam, 2015). We found that the stabilization of HIF-1α using DMOG restores the M1 functions in PE-M1 macrophages. While DMOG treatment is known to increase HIF-1α protein levels, we also observed an increase in mRNA expression. As the positive feedback loop between HIF-1α and aerobic glycolysis has been reported (Braverman *et al.*, 2016), we consider that recovering the glycolytic flux in DMOG-treated PE-M1 macrophages can increase HIF-1α expression at both transcriptional and post-transcriptional levels.

Concerning the possible mechanism by which TB-PE can impair the glycolytic pathway in M1 macrophages, it was recently established that TB-PE leads to the accumulation of lipid bodies in M-CSF-driven macrophages (Genoula *et al.*, 2018), thus changes in the lipid metabolism may have an impact on the rewiring of the metabolism of M1 macrophages. In this regard, it is important to highlight that the host-derived lipid fraction of TB-PE was enough to inhibit the aerobic glycolysis of these cells, as well as to drive the formation of foamy macrophages (data not shown). Similarly, hypercholesterolemia alters macrophage functions and metabolism by suppressing the LPS-mediated induction of the pentose phosphate pathway (Baardman *et al.*, 2018).

Despite the potential of our findings, some concerns remain to be addressed including: 1) whether TB-PE truly reflects what happens in pulmonary tissue, or whether it only represents the situation found in extra-pulmonary TB; 2) which lipids present in TB-PE are involved; and 3) timing and duration of the metabolic changes observed in macrophages in the course of the infection. Concerning the last issue, a recent study demonstrated that, unlike short-term infections, persistent infections with Mtb attenuates glycolysis in macrophages over time (Hackett *et al.*, 2020).

We demonstrated that HIF-1α stabilization improves mycobacterial control in Mtb-infected mice. DMOG promoted M1-like metabolism in tissue resident AMs, which are known to be biased towards an M2-like phenotype along with susceptibility to intracellular bacterial growth (Huang *et al.*, 2018; Pisu *et al.*, 2020). AMs from DMOG-treated animals displayed a reduction in the mitochondrial inner membrane potential and an increase in the mROS production, indicating that DMOG treatment may protect against Mtb infection associated with a metabolic switch towards glycolysis. It is also important to highlight that our results are constrained to early time points of infection (14 days). In fact, it was reported that, unlike early infection, HIF-1α inhibition diminished bacillary loads at later time points, suggesting that sustained activation of HIF-1α may result in pathological inflammation worsening the disease (Baay-Guzman *et al.*, 2018)

Recently, the regulation of glycolytic shift in metabolism by Akt-mTOR/HIF-1α signaling was elucidated with respect to trained immunity in macrophages (Cheng *et al.*, 2014). Thus, the impairment of the mTOR/HIF-1α axis by the TB-associated microenvironment may have important clinical implications leading to the long-term modulation of the innate immunity. This knowledge may contribute to future TB vaccine development by combining both innate and adaptive arms of the immune memory. Besides, several strategies aimed to modulate HIF1a activity have been implemented in pre-clinical and clinical trials *(i.e.,* targeting HIF-prolyl hydroxylases for anemia therapy (Haase, 2017) or for combating skin pathogens (Okumura *et al.*, 2012)), supporting the possible therapeutic impact of HIF modulation in disease settings.

Based on all these findings, we propose that human macrophages committed towards a pro-inflammatory profile to control the infection can be metabolically reprogrammed by host lipids generated during the Mtb infection. Reprogrammed macrophages exhibited an OXPHOS metabolic program along with attenuated pro-inflammatory and microbicide properties, resulting in susceptibility to Mtb intracellular growth.

In conclusion, together with our previous work, this study contributes to the establishment of the acellular fraction of TB-PE as a physiologically relevant tuberculous lung-milieu, demonstrating its capacity to shift macrophage metabolism to the advantage of the pathogen. It remains to be elucidated whether this metabolic shift also benefits the host as a disease tolerance mechanism to avoid tissue damage. Yet, we are certain its use will be complementary to current models of TB investigation, providing better understanding of the complex interaction between the host immune response and the bacilli, and the lung environment where it takes place.

## ACKNOWLEDGMENTS

We thank Alison Charton for her key help in the *in vivo* experiments, the TRI-IPBS platform (Emmanuelle Naser and Antonio Peixoto) for flow cytometry and image analyses, the ANEXPLO-IPBS platform (Flavie Moreau and Céline Berrone) for maintenance of ASB3 facility and BSL3 multi-pathogen facility, and the MetaToul Lipidomic Core Facility (I2MC, Inserm 1048, Toulouse, France, MetaboHUB-ANR-11-INSB-0010), for assistance and access to the mass spectrometer instrument. In addition, we are grateful for the editing service provided by Life Science editors.

This work was supported by the Argentinean National Agency of Promotion of Science and Technology (PICT-2015-0055 to MCS and PICT-2017-1317 to LB), the Argentinean National Council of Scientific and Technical Investigations (CONICET, PIP 112-2013-0100202 to MCS), and CONACYT/CONICET, the National Council for Science and Technology Mexico (CONACyT FC 2015-1/115 to MCS and CST), the Centre National de la Recherche Scientifique, the Université Toulouse-III Paul Sabatier, the Agence Nationale de la Recherche (ANR-18-CE15-0004-01 to DH), Fondation pour la Recherche Medicale (ING20160435108 to EL), and the Agence Nationale de Recherche sur le Sida et les Hépatites Virales (ANRS2014-CI-2, ANRS2014-049 to ON, and ANRS 2020-1 to CV and GL-V). The funders had no role in study design, data collection, and analysis, decision to publish, or preparation of the manuscript.

## AUTHOR CONTRIBUTIONS

Conceptualization & methodology: JLMF, DC, JPC, EL, YR, DC, DH, GL-V and LB. Investigation: JLMF, MG, DC, GD, MF, MM, MBD, OEAT, EPM, EL, GL-V, LB. Resources: EJM, DP, MO, PS, CST, RHP, JPC, DH, ON, CV, MdCS and LB. Writing: DP, MO, JPC, DH, ON, CV DC, EL, GL-V, MdCS and LB. Visualization: VS, CV and FF. Funding acquisition: DH, ON, MdCS and LB. Corresponding authors: LB is responsible for ownership and responsibility that are inherent to aspects of this study.

## DECLARATION OF INTERESTS

The authors have declared that no conflict of interest exists.

## STAR Methods

### CONTACT FOR REAGENT AND RESOURCE SHARING

Further information and requests for reagents should be directed to and will be fulfilled by the Lead Contact, Geanncarlo Lugo-Villarino. Sharing of antibodies and other reagents with academic researchers may require UBMTA agreements.

### EXPERIMENTAL MODEL AND SUBJECT DETAILS

#### Human Subjects

Buffy coats from healthy donors were prepared at Centro Regional de Hemoterapia Garrahan (Buenos Aires, Argentina) according to institutional guidelines (resolution number CEIANM-664/07). Informed consent was obtained from each donor before blood collection.

PE were obtained by therapeutic thoracentesis by physicians at the Hospital F. J Muñiz (Buenos Aires, Argentina). The diagnosis of TB pleurisy was based on a positive Ziehl-Nielsen stain or Lowestein-Jensen culture from PE and/or histopathology of pleural biopsy, and was further confirmed by an Mtb-induced IFN-γ response and an adenosine deaminase-positive test (Light, 2010). Exclusion criteria included a positive HIV test, and the presence of concurrent infectious diseases or non-infectious conditions (cancer, diabetes, or steroid therapy). None of the patients had multidrug-resistant TB. Particularly, we studied 31 patients with PE that were divided according their aetiology. First group had 12 patients with tuberculous PE; second group had 6 patients with malignant PE (PE-C) including mesothelioma, lung carcinoma or metastatic PE; third group had 8 patients with parapneumonic PE (PE-PNI); and forth group had 5 patients with transudates secondary to heart failure (HF-PE).

The research was carried out in accordance with the Declaration of Helsinki (2013) of the World Medical Association and was approved by the Ethics Committees of the Hospital F. J Muñiz and the Academia Nacional de Medicina de Buenos Aires (protocol number: NIN-1671-12, renewed in 2018; and 12554/17/X, respectively). Written informed consent was obtained before sample collection.

#### Animal Studies

Six-to eight-week-old female mice C57BL/6 were purchased from Charles River Laboratories. The experiments with these animals were performed in animal facilities based on the legal requirements in France and by qualified personnel that guarantee and minimize potential discomfort for the animals.

The procedures, including animal studies, were strictly conducted according with French laws and regulations in compliance with the European community council directive 68/609/EEC guidelines and its implementation in France.

## METHOD DETAILS

### Bacterial strain and antigens

The Mtb laboratory strain, H37Rv, was grown at 37°C in Middlebrook 7H9 medium supplemented with 10% albumin-dextrose-catalase (both from Becton Dickinson, New Jersey, USA) and 0.05% Tween-80 (Sigma-Aldrich, St. Louis, MO, USA). The Mtb γ- irradiated H37Rv strain (NR-49098) and corresponding total lipid preparation (NR-14837) were provided by BEI Resources, USA.

### Preparation of PE Pools

The PE were collected in heparin tubes and centrifuged at 300 g for 10 minutes at room temperature without brake. The cell-free supernatant was transferred into new plastic tubes, further centrifuged at 12,000 g for 10 minutes and aliquots were stored at −80°C. After having the diagnosis of the PE, pools were prepared by mixing same amounts of individual PE associated to a specific etiology. The pools were decomplemented at 56°C for 30 minutes and filtered by 0.22 μm in order to remove any remaining debris or residual bacteria.

### Preparation of Human Monocyte-Derived Macrophages

Peripheral blood mononuclear cells were obtained by Ficoll gradient separation on Ficoll-Paque (GE Healthcare, Amersham, UK). Then, monocytes were purified by centrifugation on a discontinuous Percoll gradient (GE Healthcare) as previously described (Genoula *et al.*, 2018). After that, monocytes were allowed to adhere to culture plates (Costar) for 1 h at 37°C in warm RPMI-1640 medium (ThermoFisher Scientific, Waltham, MA). The cells were then washed with warm phosphate buffered saline (PBS) twice. The final purity was checked by fluorescence-activated cell sorting analysis using an anti-CD14 monoclonal antibody (mAb) and was found to be >90%. The medium was then supplemented to a final concentration of 10% fetal bovine serum (FBS, Sigma-Aldrich) and human recombinant Granulocyte-Macrophage Colony-Stimulating Factor (GM-CSF, Peprotech, New Jork, USA) at 50 ng/ml. Cells were allowed to differentiate for 5-7 days leading to M0 macrophages.

### Macrophages treatments

Macrophages were polarized towards M1 profile with IFN-γ (10 ng/ml) plus LPS (10 ng/ml), and then exposed (or not) to 20% v/v of PE during 24 h to generate PE-M1 and M1 cells, respectively. Alternatively, macrophages were stimulated with IFNγ plus either irradiated Mtb (equivalent to a multiplicity of infection (MOI) of 2:1) or viable Mtb (MOI 5:1), and exposed (or not) to 20% v/v of PE for 24 h.

### Determination of metabolites

Lactate production and glucose concentrations in the culture medium was measured using the spectrophotometric assays Lactate Kit and Glicemia Enzimática AA Kit both from Wiener (Argentina), which are based on the oxidation of lactate or glucose, respectively, and the subsequent production of hydrogen peroxide (Trinder, 1969; Barham and Trinder, 1972). In particular, the consumption of glucose was determined by assessing the diminution of glucose levels in culture supernatants in comparison with RPMI 10% FBS. The absorbance was read using a Biochrom Asys UVM 340 Microplate Reader microplate reader and software.

### Determination of glucose uptake

Macrophages were incubated with the fluorescent glucose analog 2-(N-(7-Nitrobenz-2-oxa-1,3-diazol-4-yl)Amino)-2-Deoxyglucose (2-NBDG) (10 μM, Invitrogen, California, USA) in PBS for 30 min. Thereafter, cells were washed and intracellular 2-NBDG was measured by flow cytometry.

### Measurement of cell respiration with Seahorse flux analyzer

XFp Extracellular Flux Analyzer (Seahorse Bioscience) can measure the rate of mitochondrial oxidative phosphorylation, through determination of the real-time oxygen consumption rate (OCR) measured in picomoles/min, on a 96 microplate-based assay platform. For this methodology, basal respiration was calculated as the last measurement before addition of the electron transport chain inhibitor oligomycin (OM) at 3 μM minus the non-mitochondrial respiration, which is the minimum rate measurement after addition of electron transport chain inhibitors rotenone (ROT) and antimycin (AA) at 0.5 μM. Estimated ATP production designates the last measurement before addition of OM minus the minimum rate after OM. Maximal respiration rate (max) was defined as the OCR after addition of OM and FCCP. Spare respiration capacity (SRC) was defined as the difference between max and basal respiration. Following this approach, OCR was determined in macrophages (1.6×10^5^ cells/well) in 3 wells for each condition. The assay was performed in XF Assay Modified DMEM. Three consecutive measurements were performed under basal conditions and after the sequential addition of the following electron transport chain inhibitors.

### Measurement of cell respiration with Oroboros

Cells were harvested and centrifuged, and the cellular pellet was resuspended in RPMI-1640 medium to the desired cellular concentration (typically in a range of 0.8 - 1 × 10^7^ cells/ml). Respiration rate of intact cells was measured at 37°C with Oroboros high resolution respirometer Oxygraph-2k (Oroboros Instruments GmbH, Austria). Oxygraph-2k possesses two separate 2-mL chambers equipped with polarographic oxygen sensors that can measure in real time both the oxygen concentration (nanomoles/milliliter) and the oxygen consumption (picomoles/second/milliliter) within each chamber. The basic principle of the system is to measure the concentration and consumption of oxygen by injecting substrates directly to cells that are suspended in solution within the chamber. In our case, routine respiration was obtained soon after succinate (10mM) addition to the system. Thereafter, OM (10mM) was added to inhibit OXPHOS respiration, allowing the measurement of the contribution of mitochondrial respiration leak. By using ROT (1mM) and AA (4mM), we determined the residual oxygen consumption, ROX, which is respiration due to oxidative side reactions remaining after inhibition of the electron transfer-pathway. These ROX determinations were used to correct the values of ORC associated to routine and leak. With these parameters, we then calculated OXPHOS activity as follows: OXPHOS = (Routine - ROX) - (Leak - ROX). To determine the activity of complex IV of the electron transfer-pathway, ascorbate (ASC, 5mM) and *N,N,N’N’,* tetramethyl-*p*-phenylenediamine (TMPD, 5mM), artificial electron donors reducing cytochrome *c*, were added to obtain the maximal activity associated to complex IV, which was then inhibited by adding potassium cyanide (KCN, 25mM). The values of ORC before and after KCN were used to calculate the net activity of complex IV (CIV). Data was analyzed with DatLab 4 software (Oroboros Instruments, Austria).

### Phenotypic characterization by flow cytometry

Macrophages were centrifuged for 7 minutes at 1200 rpm and then stained for 40 minutes at 4°C with fluorophore-conjugated antibodies PE-anti-Glut1 (clone 202915 R&D Systems, Minnesota, USA) and in parallel, with the corresponding isotype control antibody. Glut1 expression was also confirmed by labelling the cells with the ligand for Glut1, H2RBD-EGFP (Kinet *et al.*, 2007). After staining, the cells were washed with PBS 1X, centrifuged and analyzed by flow cytometry using FACSCalibur cytometer (BD Biosciences, San Jose, CA, USA). For HIF-1α and mTOR determination, macrophages were permeabilized with either methanol for HIF-1α or Perm2 solution for mTOR (BD Biosciences), and incubated with PE-anti-HIF-1α (clone 546-16, Biolegend, San Diego, USA) or PE-anti-mTOR (pS2448, BD Biosciences), respectively. The macrophage population was gated according to its Forward and Size Scatter properties. The median fluorescence intensity (MFI) was analyzed using FCS Express V3 software (De Novo Software, Los Angeles, CA, USA). For mice experiments, cell surface staining of single-cell suspensions from lungs was performed using fluorophore-conjugated antibodies: CD45.2, MERTK (clone DS5MMER), SIGLEC-F (clone E50-2440), CD64 (clone 10.1), Ly6G (RB6-8C5), CD11c (clone N418), CD11b (clone M1/70) and Live/Dead (LIVE/DEAD™ Fixable Aqua Dead Cell, Thermofisher Scientific). For analysis of mitochondrial markers, cells were stained with MitoSOX Red (5μM) and DiOC6 (40nM) (ThermoFisher) at 37°C for 30 min. Cell staining was analyzed using a FACSAria Fusion cytometer in BSL3 facility and FlowJo software version V10. Cells were first gated on singlets (FSC-H vs. FSC-W and SSC-H vs. SSC-W) and live cells before further analyses.

### Quantitative RT-PCR

Total RNA was extracted with Trizol reagent (Thermo Fisher Scientific) and cDNA was reverse transcribed using the Moloney murine leukemia virus reverse transcriptase and random hexamer oligonucleotides for priming (Life Technologies, CA, USA). The expression of the genes of hexokinase 2 (HK2) and LDH-A was determined using PCR SYBR Green sequence detection system (Eurogentec, Seraing, Belgium) and the CFX Connect™ Real-Time PCR Detection System (Bio-Rad, CA, United States). Gene transcript numbers were standardized and adjusted relative to eukaryotic translation elongation factor 1 alpha 1 (EeF1A1) transcripts. Gene expression was quantified using the ΔΔCt method. Primers used for RT-PCR were as follows: EeF1A1 Fwd: 5’-CCAAGACCCAGGCATACTTGGA-3’ and Rev: 5’-TCGGGCAAGTCCACCACTAC-3’; HK2 Fwd: 5’-GAGCCACCACTCACCCTACT-3’ and Rev: 5’-CCAGGCATTCGGCAATGTG-3’; and LDH-A Fwd: 5’-TGGGAGTTCACCCATTAAGC-3’ and Rev: 5’-AGCACTCTCAACCACCTGCT-3’.

### Fluorescence microscopy

Cells were stained with 250 nM MitoSpy Green FM (Biolegend) for 30 minutes. The cells were then washed with PBS three times. All the samples were imaged on a FluoView FV1000 confocal microscope (Olympus, Tokyo, Japan) equipped with a Plapon 60X/NA1.42 objective. To quantify the mitochondrial morphology of MitoSpy Green FM-stained macrophages, we developed a computer-aided analysis tool scoring the mitochondrial length and number per cell with ImageJ-Fiji software. Briefly, we used a motorized z-focus to obtain 3D images spanning the entire height of blindly selected cells to capture the entire architecture of the mitochondrial network and combined them into a single image using a maximum intensity projection. A threshold was defined to identify mitochondria based on their fluorescence intensity, and images were ‘skeletonized’ to measure mitochondrial length (perimeter) and number. At least 20 cells per condition were analyzed in each independent experiment.

### Transmission electron microscopy

Macrophages were fixed in 2.5% glutaraldehyde/ 2% paraformaldehyde (EMS, Delta-Microscopies) dissolved in 0.1 M Sorensen buffer (pH 7.2) during 2h at room temperature, and then preserved in 1 % paraformaldehyde (PFA) dissolved in Sorensen buffer. Adherent cells were treated for 1 h with 1% aqueous uranyl acetate then dehydrated in a graded ethanol series and embedded in Epon. Sections were cut on a Leica Ultracut microtome and ultrathin sections were mounted on 200 mesh onto Formvar carbon-coated copper grids. Finally, thin sections were stained with 1% uranyl acetate and lead citrate and examined with a transmission electron microscope (Jeol JEM-1400) at 80 kV. Images were acquired using a digital camera (Gatan Orius).

### Changes of mitochondrial membrane potential

Mitochondrial membrane potential was measured using the mitochondrial-specific dual-fluorescence probe, 5,5’,6,6’-tetrachloro-1,1 ‘,3,3’-tetraethylbenzimidazolylcarbocyanine iodide (JC-1, Molecular Probes, Eugene, OR, USA). Macrophages were washed twice in ice-cold PBS and loaded with 2 μM JC-1 for 30 min, and then measured by flow cytometry (FACScan, BD Biosciences). In parallel, mitochondrial mass was determined in individual cells by labelling them with the probe MitoSpy Green FM (Biolegend). Green fluorescence was analyzed by flow cytometry.

### Reactive oxygen species measurements

To measure the mROS superoxide, macrophages were incubated with MitoSOX (Molecular Probes) at 2.5 μM in serum-free RPMI media for 15 to 30 minutes at 37°C. Cells were washed with warmed PBS, removed from plates with cold PBS containing 1 mM ethylenediamine tetraacetic acid (EDTA) by pipeting, pelleted at 1500 rpm for 3 minutes, immediately resuspended in cold PBS containing 1% FBS, and subjected to FACS analysis.

### Soluble II.-1β determinations

The amounts of human IL-1β were measured by ELISA, according to manufacturer’s instructions kits (ELISA MAX™ Deluxe Kits from Biolegend). The detection limit was 15.6 pg/ml.

### Western blots

Macrophages were lysed in ice-cold buffer consisting of 150 mM NaCl, 10 mM Tris, 5 mM EDTA, 1% Sodium Dodecyl Sulfate (SDS), 1% Triton X-100, 1% sodium deoxycholate, gentamicin/streptomycin, 0.2% azide plus a cocktail of protease inhibitors (Sigma-Aldrich). Lysates were incubated on ice for 3 h and cleared by centrifugation for 15 minutes at 14,000 rpm at 4°C. Protein concentrations were determined using the bicinchoninic acid (BCA) protein assay (Pierce, ThermoFisher). Equal amounts of protein (40 μg) were then resolved on a 10% SDS-PAGE. Proteins were transferred to Hybond-ECL nitrocellulose membranes (GE Healthcare) for 2 h at 100 V and blocked with 1% Bovine Serum Albumine (BSA)-0.05% Tween-20 for 1 h at room temperature. Membranes were probed with primary anti-human nuclear factor-kappa B (NF-κB) p65 (c-20) sc-372 (1:200 dilution, Santa Cruz Biotechnology, Palo Alto, CA, USA) overnight at 4°C. After extensive washing, blots were incubated with a horseradish peroxidase (HRP)-conjugated goat anti-rabbit IgG Ab (1:5000 dilution; Santa Cruz Biotechnology) for 1 h at room temperature. Immunoreactivity was detected using ECL Western Blotting Substrate (Pierce). Protein bands were visualized using Kodak Medical X-Ray General Purpose Film. For internal loading controls, membranes were stripped by incubating in buffer consisting of 1.5% Glycine, 0.1% SDS, 1% Tween-20, pH 2.2 for 10 minutes twice, extensively washed and reprobed with anti-β-actin (1:2000 dilution; ThermoFisher, clone AC-15) and or HRP-conjugated goat antimouse IgG Ab (1:2000 dilution; Santa Cruz Biotechnology). Results from Western blot were analyzed by densitometric analysis (Image J software).

### Infection of human macrophages with Mtb

Infections were performed in the biosafety level 3 (BSL-3) laboratory at the *Unidad Operativa Centro de Contención Biológica (UOCCB), ANLIS-MALBRAN* (Buenos Aires), according to the biosafety institutional guidelines. Macrophages were infected with Mtb H37Rv strain at a MOI of 1-5:1 during 1 h at 37°C. Next, extracellular bacteria were removed gently by washing with pre-warmed PBS, and cells were cultured in RPMI-1640 medium supplemented with 10 % FBS and gentamicin (50 μg/ml).

### Measurement of bacterial intracellular growth in macrophages by colony forming units (CFU) assay

Macrophages stimulated or not with IFN-γ were infected with Mtb H37Rv strain at a MOI of 0.5 bacteria/cell in the presence (or not) of TB-PE (triplicates). After 24 h, extracellular bacteria were removed by gently washing four times with pre-warmed PBS. At days 3 and 6 cells were lysed in 0.1 % SDS and neutralized with 20 % BSA in Middlebrook 7H9 broth. Serial dilutions of the lysates were plated in triplicate, onto 7H11-Oleic Albumin Dextrose Catalase (OADC, Becton Dickinson) agar medium for CFU scoring at 21 days later.

### Visualization and quantification of Mtb infection

Macrophages seeded on glass coverslips within a 24-well tissue culture plate at a density of 5 × 10^5^ cells/ml were infected with the red fluorescent protein (RFP) expressing Mtb CDC 1551 strain at a MOI of 5:1 during 2 h at 37°C. Then, extracellular bacteria were removed gently by washing with pre-warmed PBS, and cells were cultured in RPMI-1640 medium supplemented with 10% FBS for 48 h. The glass coverslips were fixed with PFA 4% and stained with BODIPY 493/503 (Life Technologies). Finally, slides were mounted and visualized with a FluoView FV1000 confocal microscope (Olympus, Tokyo, Japan) equipped with a Plapon 60X/NA1.42 objective, and analyzed with the software ImageJ-Fiji. We measured the occupied area with RFP-Mtb (expressed as Raw Integrated Density) per cell in z-stacks from confocal laser scanning microscopy images. Individual cells were defined by boron-dipyrromethene (BODIPY)-stained cellular membranes, which allow us to define the region of interests for quantification. For quantification 80-100 cells of random fields per condition were analyzed.

### Mtb H37Rv infection and treatment with DMOG of mice

Experimental mice were anesthetized with a cocktail of ketamine (60 mg/kg, Merial) and xylasine (10 mg/kg, Bayer) and infected intranasally with ≈1000 CFUs of the Mtb experimental strain, H37Rv, in 25 μL of PBS. For drug treatment, mice received 1.25 mg of DMOG (Bertin Technologies) intraperitoneally per mouse every day starting upon infection for 14 days.

After the mice were sacrificed, the lungs were harvested aseptically, homogenized using a gentle MACS dissociator (C Tubes, Miltenyi, Biotec, CA, USA), and incubated with DNAse I (0.1 mg/mL, Roche) and collagenase D (2 mg/mL, Roche) during 30 minutes at 37°C under 5% CO2. The Mtb load was determined by plating serial dilutions of the lung homogenates onto 7H11 solid medium supplemented with OADC (Middlebrook). The plates were incubated at 37°C for 3 weeks before bacterial CFUs scoring. Lungs homogenates were filtered on 40 μm cell strainers and centrifuged at 329 × *g* during 5 min. Red blood cells were lysed in 150 mM NH_4_Cl, 10 mM KHCO_3_, 0.1 mM EDTA (pH 7.2).

### Total lipid and polar metabolite extractions

Polar metabolites (PMPE) and lipids (LPE) extractions from TB-PE were based on Bligh and Dyer protocol. Briefly, 1 ml of TB-PE was transferred to 5 ml glass tubes containing 1.5 mL CHCl3/CH3OH (2:1, v/v). The samples were incubated overnight at 4°C with gentle agitation. After centrifugation, top layers were subjected to an additional extraction using CHCl_3_:CH_3_OH (1:2, v/v) and centrifuging at high speed for 2 minutes. The upper aqueous and organic lower phases, primarily PMPE and LPE, respectively, were collected and dried under vacuum. PMPE were resuspended in 12.5 μl of water, while LPE in 12.5 μl of DMSO. PMPE and LPE fractions were used at the same concentration as in the untouched TB-PE (1x) or at a double dose (2x).

### Lipidomic analysis

For the screening of the *M. tuberculosis* lipid families, TB-PE lipid extracts were resuspended at 0.5 mg/mL in CHCl_3_/CH_3_OH (1:1, v/v) and analyzed using a Waters Xevo G2-XS Q-Tof mass spectrometer coupled to a Waters Supercritical Fluid Chromatography ACQUITY UPC2 at the MetaToul Lipidomic Core Facility (I2MC, Inserm 1048, Toulouse, France). One microliter of lipid extract was injected on a Waters Torus DIOL column (1.7 μm x 100 mm x 3 mm) and the lipid families were separated using a gradient of CH_3_OH (1-50 %) in CO_2_, which allows the detection of Mtb lipids families with characteristic retention time, acylform profiles and accurate m/z, as previously described (Layre *et al.*, 2011). Spectra were collected in positive and negative-ion mode from m/z 100 to 3000. Characteristic signals of phthiocerol dimycocerosate, glucose/trehalose monomycolate, sulfoglycolipids, tuberculosinyl adenosines, triglycosylated phenolic glycolipids or phosphatidyl-*myo*-inositol mannosides, could not be extracted from the dataset obtained for LPE samples (within 10 ppm accuracy).

## QUANTIFICATION AND STATISTICAL ANALYSIS

All values are presented as mean and standard error of the mean SEM of a number of independent experiments. Independent experiments are defined as those performed with macrophages derived from monocytes isolated independently from different donors. As most of our datasets did not pass the normality tests, non-parametric tests were applied. Comparisons between more than two paired data sets were made using the Friedman test followed by Dunn’s Multiple Comparison Test. Comparisons between two paired experimental conditions were made using the two-tailed Wilcoxon Signed Rank. For the analysis of the OCR measurements, t-test was applied. Comparisons between control or DMOG-treated mice were performed using the Mann-Whitney test. For all statistical comparisons, a p value <0.05 was considered significant.

## Supplemental Information titles and legends

**Figure S1. TB-PE did not induce cell death in human macrophages under the implemented experimental settings. *Related to Figure 1*.** Macrophages were polarized towards M1 profile with IFN-γ (10 ng/ml) plus LPS (10 ng/ml) and were exposed or not to 20% v/v of PE during 24 h to generate PE-M1 and M1 cells respectively. SFB-deprived macrophages were used as positive controls. **(A)** Raw FACS plots illustrating the FSC/SSC features of the macrophage populations. **(B)** Lactate dehydrogenase (LDH) secretion. N=4. **(C)** Representative dot plot from 1 independent experiment of FITC-Annexin V (AV) and propidium iodide (PI) staining in macrophages. Percentages of early apoptotic (AV^+^/PI^-^), late apoptotic (AV^+^/PI^+^) or necrotic (AV^-^/PI^+^) cells. Data were obtained from 4 independent experiments. Friedman test followed by Dunn’s Multiple Comparison Test: *, p < 0.05; only the differences vs M0 are shown. **(D)** Mean fluorescence intensity (MFI) of HLA-DR, CD86, PD-L1, and CD80 measured by flow cytometry, N = 6. Representative histograms are shown. Friedman test followed by Dunn’s Multiple Comparison Test: *p < 0.05; **p < 0.01 as depicted by lines.

**Figure S2. Different Mtb strains drive the metabolic reprogramming in IFN-γ-activated macrophages. *Related to Figure 4*.** M(IFN-γ)-activated macrophages were infected with the following Mtb strains: H37Rv, H37Ra, LAM9005186, or Bei 583, at MOIs 1:1 or 5:1 Lactate release **(A)** and glucose uptake **(B)** measured in supernatant media collected at for 4 and 24h post-infection. N=4. **(C)** Raw FACS plots illustrating the FSC/SSC features of the macrophage populations. **(D)** Mean fluorescence intensity (MFI) of HLA-DR, CD86, PDL-1, and CD80 in response to irradiated Mtb measured by flow cytometry, N = 6. Representative histograms are shown. Friedman test followed by Dunn’s Multiple Comparison Test: *p < 0.05; **p < 0.01 as depicted by lines.

**Figure S3. Characterization of pulmonary phagocyte subsets from DMOG-treated mice during Mtb infection. *Related to Figure 6*.** Lungs was harvested from Ctl and DMOG-treated mice 2 wk after infection by intranasal challenge with 10^3^ Mtb. The phagocyte subsets were analyzed by flow cytometry to determine their identity. **(A)** Gate strategy to distinguish polymorphonuclear leukocytes (PMN), interstitial macrophages (IM) and, alveolar macrophages (AM). **(B)** Percentages (%) of PMN among CD45.2+ Live cells and **(C)** absolute number (#) of PMN recovered from lungs of Ctl and DMOG-treated Mtb-infected animals. A representative experiment of 2 is shown. **(D)** Representative histogram of DiOC6 labeling for IM from DMOG-treated mice, control mice and fluorescent minus one (FMO) (**E**) Mean fluorescent intensity (MFI) quantification of DiOC6 for IM (**F**) Percentages of DiOC6^high^ IM. A representative of two independent experiments is shown. (**G**) Representative histogram of MitoSOX labeling for IM from DMOG-treated mice, control mice and FMO. (**H**) MFI quantification of MitoSOX for AM. (**I**) Percentages of MitoSOX^high^ IM. A representative of two independent experiments is shown. A representative of two independent experiments is shown. Mann-Whitney test: *p < 0,05; **p< 0,01; Two-way ANOVA: *p < 0,05.

**Figure S4. Elucidation of the nature of the TB-PE components responsible for the metabolic alterations observed in human macrophages. *Related to Figure 7*. (A)** Commassie blue staining of tuberculous PE treated with polyacrylamide hydrogel-embedded Proteinase K (Pk PE) or with polyacrilamide alone (Mock PE). **(B-C)** Lactate release (B) and cell viability (C) of M0, M1, PE-M1, Mock PE-M1 and Pk PE-M1 cells, N = 10. **(D)** Raw FACS plots illustrating the FSC/SSC features of M0, M1, PE-M1, PMPE-M1 (Polar metabolites PE-M1) or LPE-M1 (Lipids PE-M1) cells. **(E)** Cell viability of macrophages exposed or not to total lipids fraction obtained from Mtb, N = 6.

## Key Resources Table

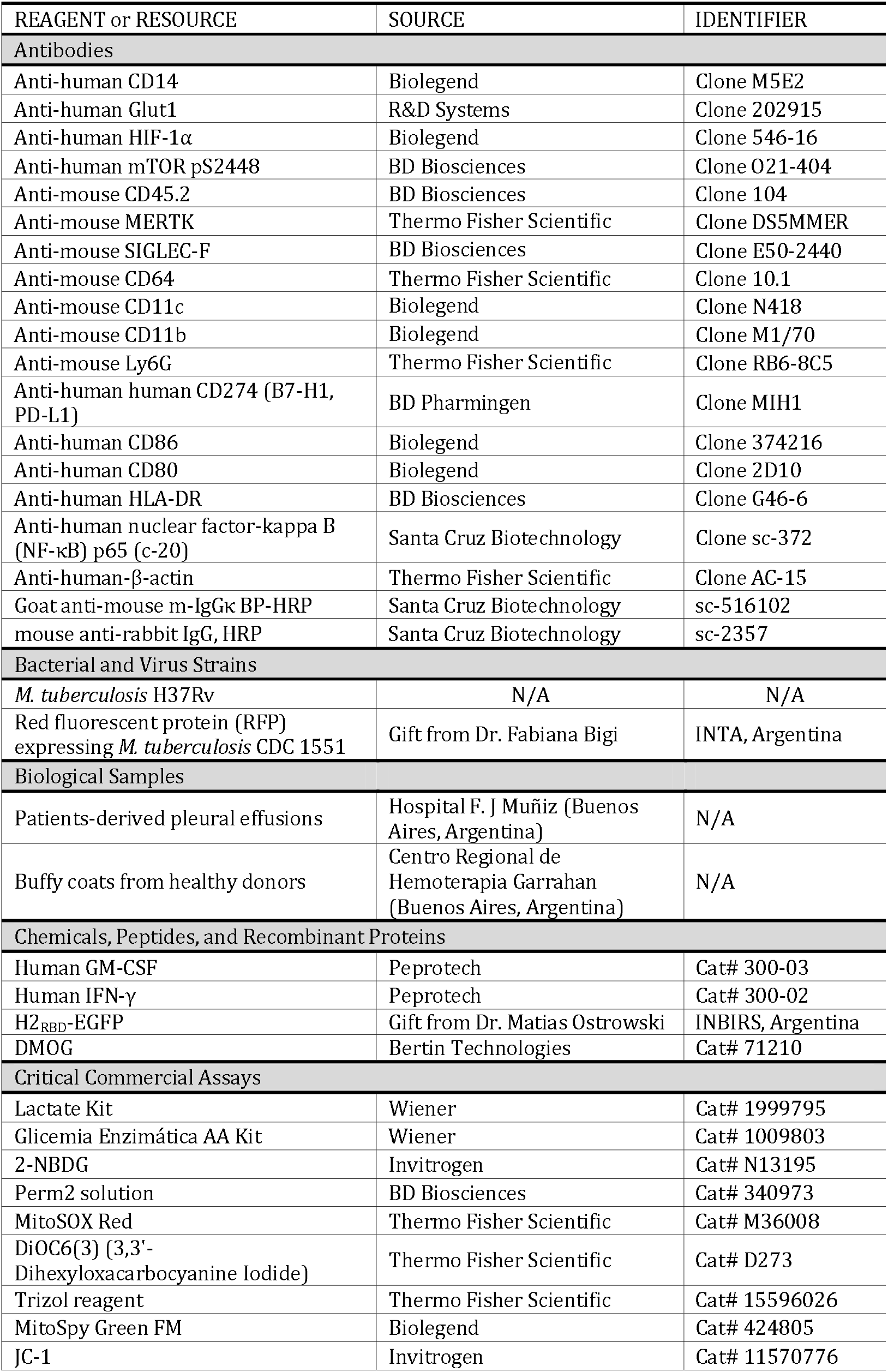
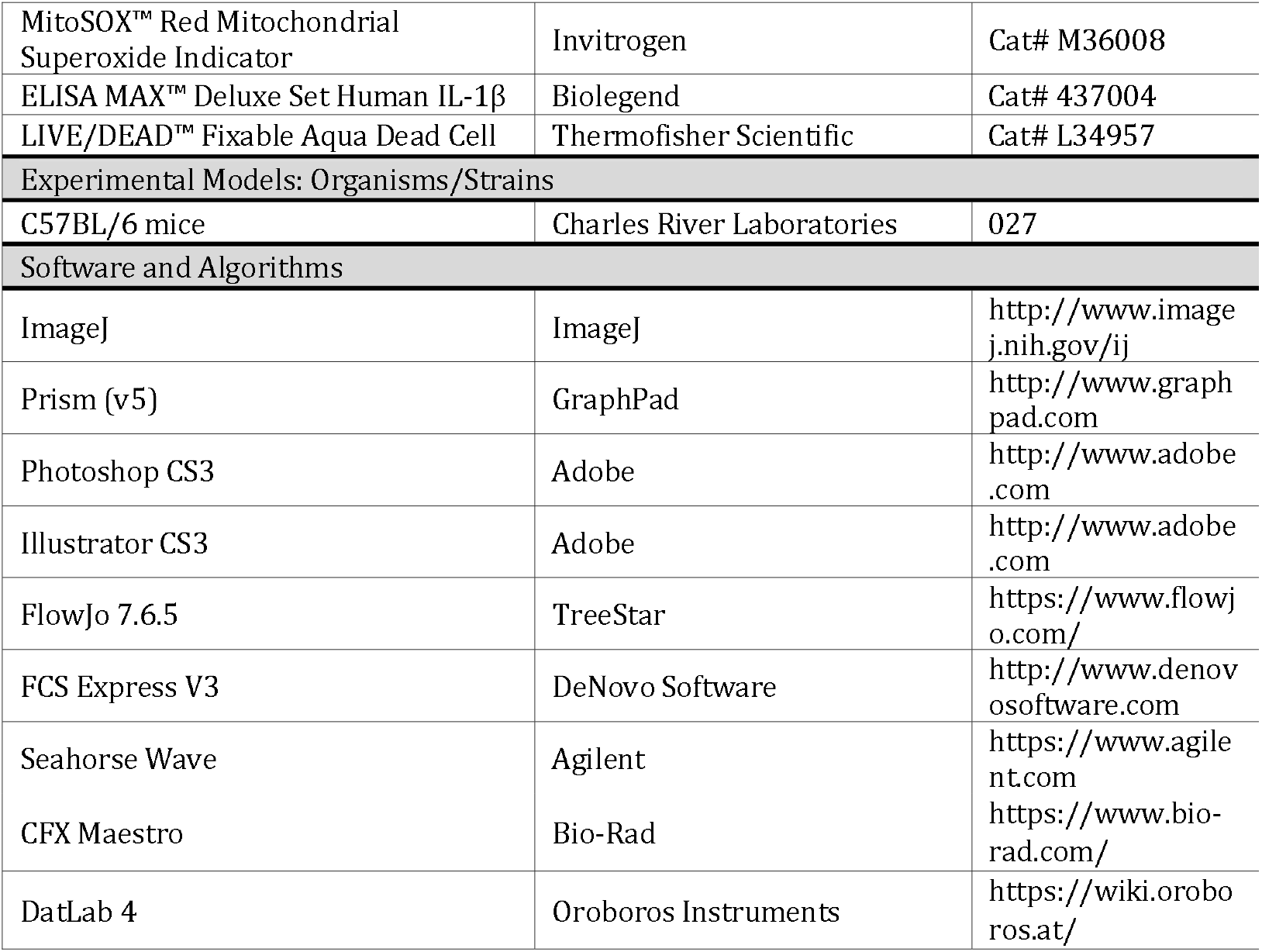

