## Supplementary Information for "Host-derived lipids from tuberculous pleurisy impair macrophage microbicidal-associated metabolic activity"

**Supplementary Material and Methods**

**Cell viability** **in macrophages. *Related to Figure S1*.** The binding of FITC-Annexin V and propidium iodide (PI) staining in macrophages were measured according to the manufacturer’s instructions (Molecular Probes, Oregon, USA), and cells were immediately analyzed by ﬂow cytometry. Cell viability was also ascertained by lactate dehydrogenase (LDH) secretion test with a kinetic UV method kit according to the manufacturer’s instructions (Roche Diagnostics GmbH, Germany).

**Phenotypic characterization by flow cytometry. *Related to Figure S1-2.*** Macrophages were centrifuged for 7 min at 1200 rpm and then stained for 40 min at 4ºC with fluorophore-conjugated antibodies FITC-anti-HLA-DR (clone G46-6, BD Biosciences), PerCP.Cy5.5-anti-CD86 (clone 374216, Biolegend), PE-anti-PD-L1 (clone MIH1, BD Pharmingen), or PE-anti-CD80 (clone 2D10, Biolegend), and in parallel, with the corresponding isotype control antibody. After staining, the cells were washed with PBS 1X, centrifuged and analyzed by flow cytometry using FACSCalibur cytometer (BD Biosciences). The macrophage population was gated according to its Forward and Size Scatter properties. The median fluorescence intensity (MFI) was analyzed using FCS Express V3 software (De Novo Software, Los Angeles, CA, USA).

**Infection of human macrophages with different Mtb strains. *Related to Figure S2*.** Macrophages were infected with the reference virulent or avirulent strains, H37Rv or H37Ra respectively, or with clinical isolates from the Latin AmericanMediterranean genotype (LAM9005186, considered a hypervirulent isolate), or the Beijing lineage (Bei 583, considered highly virulent), during 1 h at 37ºC. Then, extracellular bacteria were removed gently by washing with pre-warmed PBS, and cells were cultured in RPMI-1640 medium supplemented with 10 % FBS and gentamicin (50 µg/ml) for 4 and 24h. All strains were grown in Middlebrook 7H9 supplemented with 10% OADC. Strains were grown to logarithmic phase and measurement of the optical density determined the culture concentrations by spectrometry at 600 nm. Working aliquots were stored at -80ºC until their use. Bacillary viability was tested by measuring the number of colonies forming units (CFU).

**Proteins digestion of TB-PE**. ***Related to Figure S4*.** Proteinase K (50 ug/mL, Sigma Aldrich) was immobilized by entrapment in polyacrylamide gel, and pieces of it were incubated with TB-PE overnight at 37 ºC to generate Pk PE. As control TB-PE was treated with polyacrilamide alone (Mock PE). Pk PE and Mock PE were centrifuged at high speed and the supernatant was collected and stored at -80 ºC until used. Protein digestion was controlled using sodium dodecyl sulfate polyacrylamide gel electrophoresis (SDS-PAGE), followed by Commassie blue staining.

**Supplementary Figures**

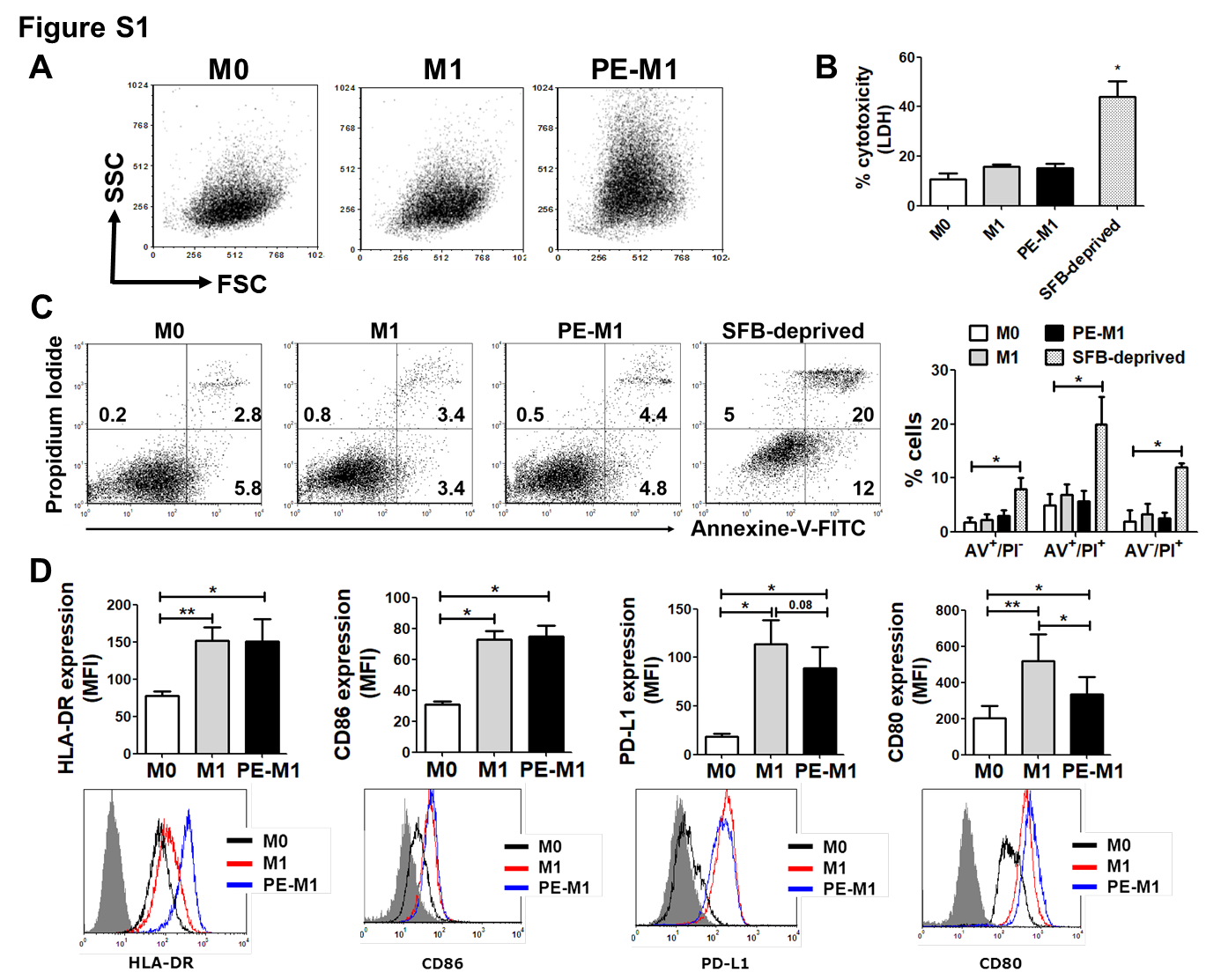

**Figure S1. TB-PE did not induce cell death in human macrophages under the implemented experimental settings. *Related to Figure 1*.** Macrophages were polarized towards M1 profile with IFN-γ (10 ng/ml) plus LPS (10 ng/ml) and were exposed or not to 20% v/v of PE during 24 h to generate PE-M1 and M1 cells respectively. SFB-deprived macrophages were used as positive controls. **(A)** Raw FACS plots illustrating the FSC/SSC features of the macrophage populations. **(B)** Lactate dehydrogenase (LDH) secretion. N=4. **(C)** Representative dot plot from 1 independent experiment of FITC-Annexin V (AV) and propidium iodide (PI) staining in macrophages. Percentages of early apoptotic (AV^+^/PI^-^), late apoptotic (AV^+^/PI^+^) or necrotic (AV^-^/PI^+^) cells. Data were obtained from 4 independent experiments. Friedman test followed by Dunn's Multiple Comparison Test: *, p < 0.05; only the differences vs M0 are shown. **(D)** Mean fluorescence intensity (MFI) of HLA-DR, CD86, PD-L1, and CD80 measured by flow cytometry, N = 6. Representative histograms are shown. Friedman test followed by Dunn’s Multiple Comparison Test: *p < 0.05; **p < 0.01 as depicted by lines.

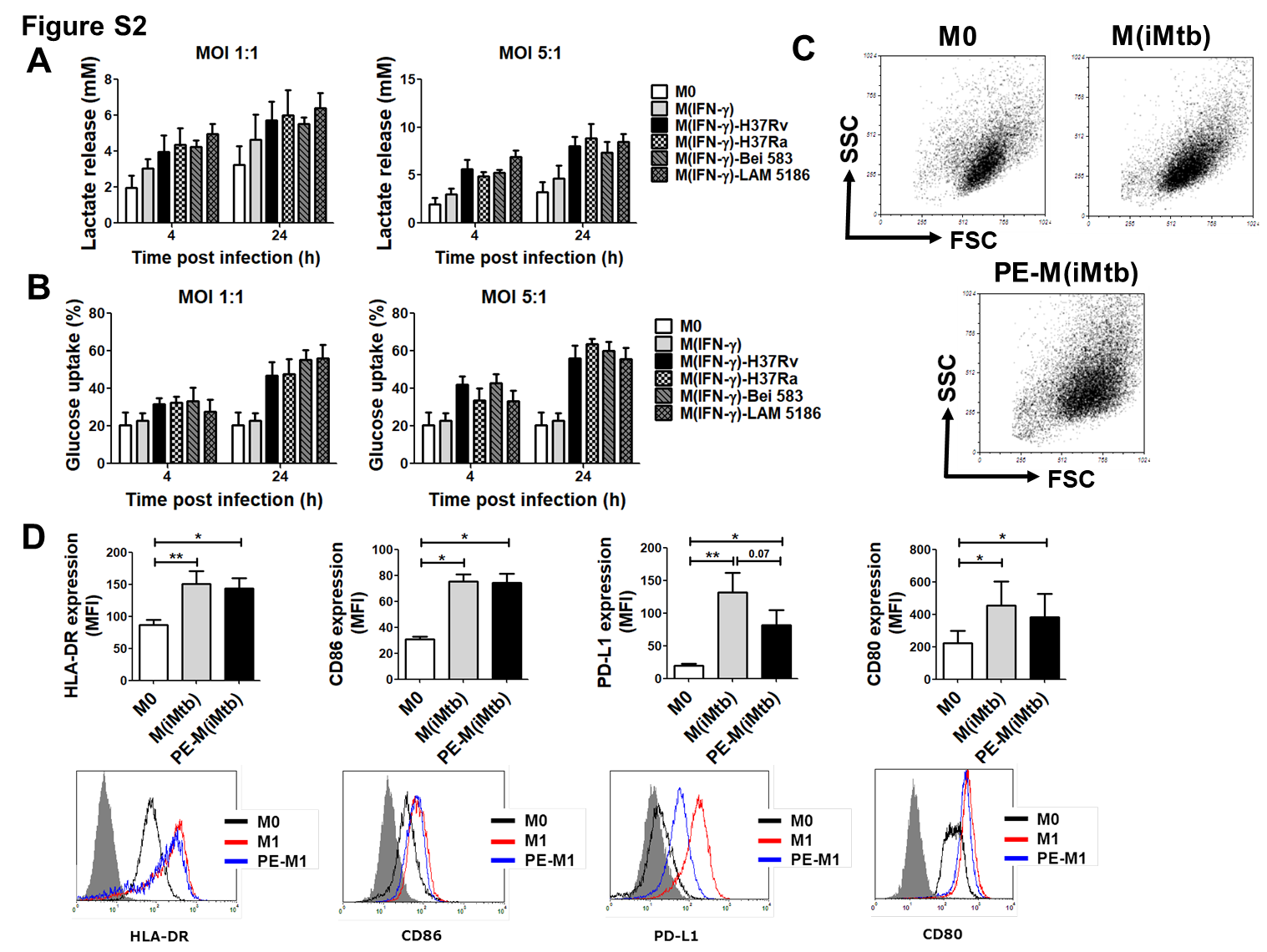

**Figure S2. Different Mtb strains drive the metabolic reprogramming in IFN-γ-activated macrophages. *Related to Figure 4.*** M(IFN-γ)-activated macrophages were infected with the following Mtb strains: H37Rv, H37Ra, LAM9005186, or Bei 583, at MOIs 1:1 or 5:1 Lactate release **(A)** and glucose uptake **(B)** measured in supernatant media collected at for 4 and 24h post-infection. N=4. **(C)** Raw FACS plots illustrating the FSC/SSC features of the macrophage populations. **(D)** Mean fluorescence intensity (MFI) of HLA-DR, CD86, PDL-1, and CD80 in response to irradiated Mtb measured by flow cytometry, N = 6. Representative histograms are shown. Friedman test followed by Dunn’s Multiple Comparison Test: *p < 0.05; **p < 0.01 as depicted by lines.

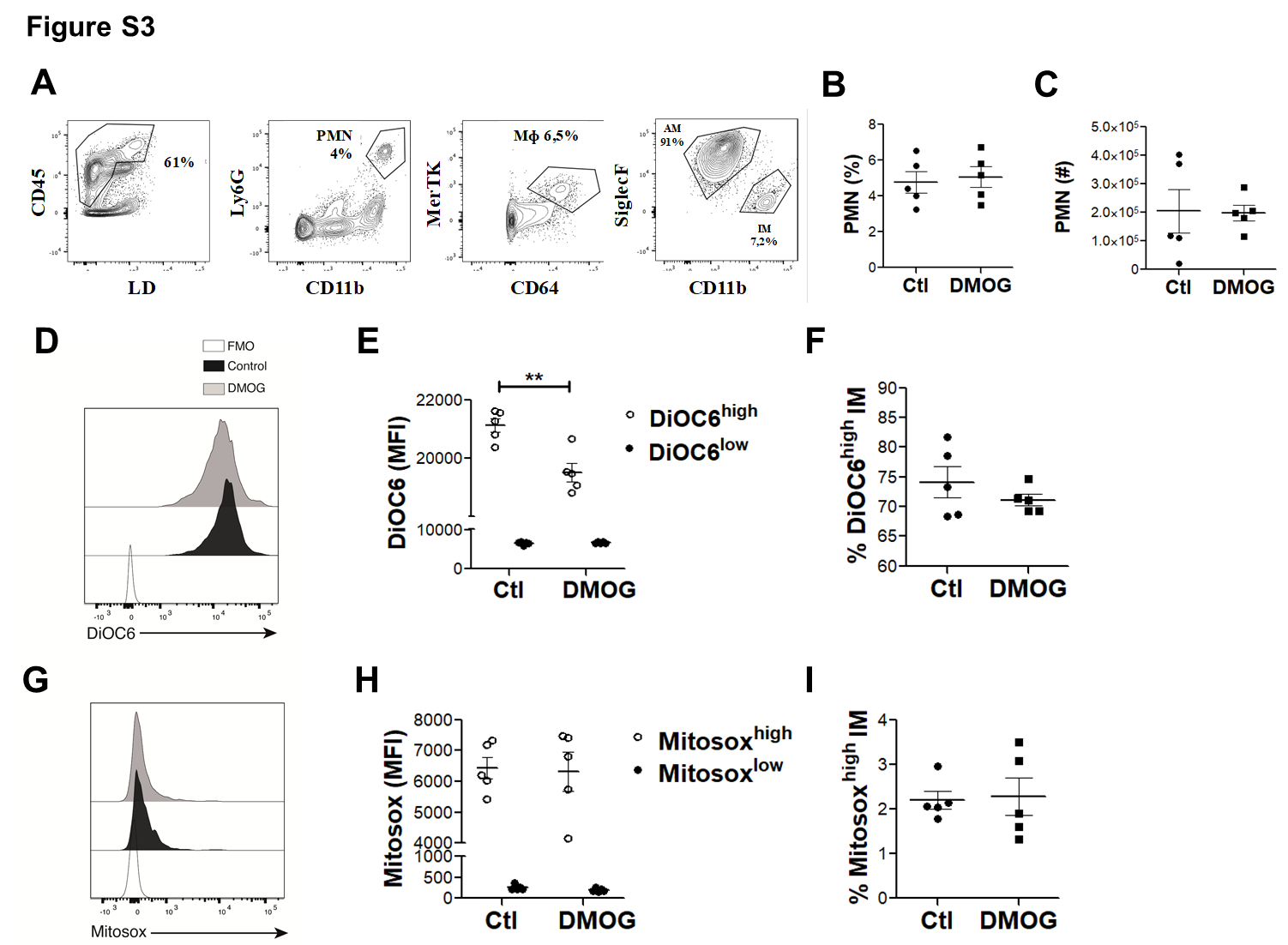

**Figure S3. Characterization of pulmonary phagocyte subsets from DMOG-treated mice during Mtb infection. *Related to Figure 6.*** Lungs was harvested from Ctl and DMOG-treated mice 2 wk after infection by intranasal challenge with 10^3^ Mtb. The phagocyte subsets were analyzed by flow cytometry to determine their identity. **(A)** Gate strategy to distinguish polymorphonuclear leukocytes (PMN), interstitial macrophages (IM) and, alveolar macrophages (AM). **(B)** Percentages (%) of PMN among CD45.2+ Live cells and **(C)** absolute number (#) of PMN recovered from lungs of Ctl and DMOG-treated Mtb-infected animals. A representative experiment of 2 is shown. **(D)** Representative histogram of DiOC6 labeling for IM from DMOG-treated mice, control mice and fluorescent minus one (FMO) (**E**) Mean fluorescent intensity (MFI) quantification of DiOC6 for IM (**F**) Percentages of DiOC6^high^ IM. A representative of two independent experiments is shown. (**G**) Representative histogram of MitoSOX labeling for IM from DMOG-treated mice, control mice and FMO. (**H**) MFI quantification of MitoSOX for AM. (**I**) Percentages of MitoSOX^high^ IM. A representative of two independent experiments is shown. A representative of two independent experiments is shown. Mann-Whitney test: *p < 0,05; **p< 0,01; Two-way ANOVA: *p < 0,05.

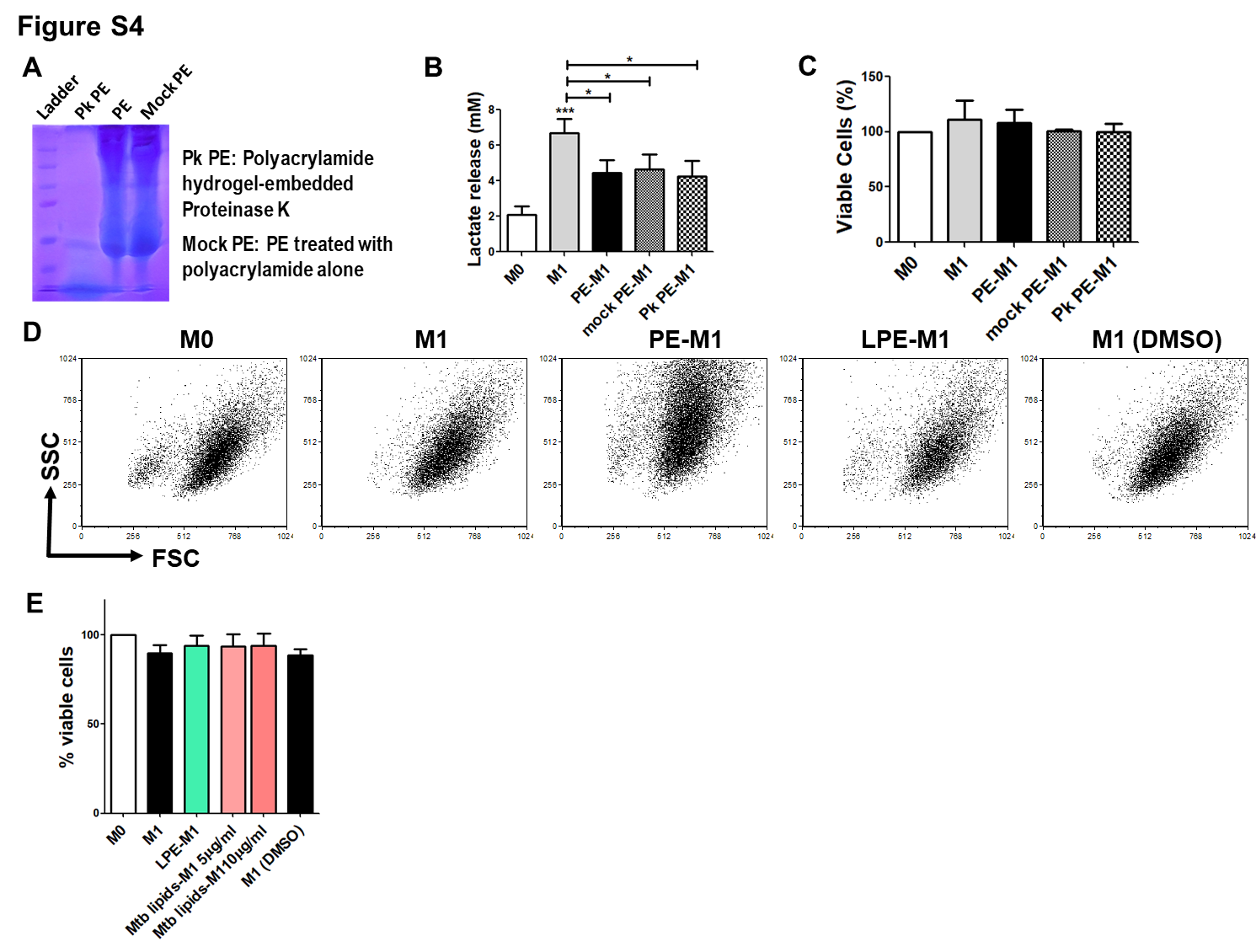

**Figure S4. Elucidation of the nature of the TB-PE components responsible for the metabolic alterations observed in human macrophages. *Related to Figure 7.* (A)** Commassie blue staining of tuberculous PE treated with polyacrylamide hydrogel-embedded Proteinase K (Pk PE) or with polyacrilamide alone (Mock PE). **(B-C)** Lactate release (B) and cell viability (C) of M0, M1, PE-M1, Mock PE-M1 and Pk PE-M1 cells, N = 10. **(D)** Raw FACS plots illustrating the FSC/SSC features of M0, M1, PE-M1, PMPE-M1 (Polar metabolites PE-M1) or LPE-M1 (Lipids PE-M1) cells. **(E)** Cell viability of macrophages exposed or not to total lipids fraction obtained from Mtb, N = 6.
